# Suppressor of Fused regulation of Hedgehog Signaling is Required for Proper Astrocyte Differentiation

**DOI:** 10.1101/2021.08.24.457497

**Authors:** Danielle M. Spice, Joshua Dierolf, Gregory M. Kelly

## Abstract

Hedgehog signaling is essential for vertebrate development, however, less is known about the negative regulators that influence this pathway. Using the mouse P19 embryonal carcinoma cell model, Suppressor of Fused (SUFU), a negative regulator of the Hedgehog pathway, was investigated during retinoic acid-induced neural differentiation. We found Hedgehog signaling was activated in the early phase of differentiation but was inactive during terminal differentiation of neurons and astrocytes. Early activation was required for neural differentiation however, it alone was not sufficient to induce neural lineages. SUFU, which regulates signaling at the level of GLI, remained relatively unchanged during differentiation, but its loss through CRISPR-Cas9 gene editing resulted in ectopic expression of Hedgehog target genes. Interestingly, these SUFU-deficient cells were unable to differentiate without retinoic acid, and when used they showed delayed and decreased astrocyte differentiation; neuron differentiation was unaffected. Ectopic activation of Hh target genes in SUFU-deficient cells remained throughout retinoic acid-induced differentiation and this was accompanied by the loss of GLI3, despite the presence of the *Gli3* message. Thus, the study indicates the proper timing and proportion of astrocyte differentiation requires SUFU, and its normal regulation of GLI3 to maintain Hh signaling in an inactive state.

## 1 Introduction

Hedgehog (Hh) signaling plays pivotal roles in neural development, where Sonic Hedgehog (SHH) is essential in patterning the differentiation of motor neurons and interneurons in the neural tube [1] and inducing differentiation of cerebellar neurons and glial cells [2]. When Hh ligand is absent, the transmembrane receptor Patched (PTCH) inhibits the transmembrane protein Smoothened (SMO) from translocating to the plasma membrane of the primary cilium [3]. Inhibition allows the Suppressor of Fused (SUFU) to sequester full length GLI transcription factors [4] or promotes GLI phosphorylation [5] and partial proteosomal degradation into a truncated repressor [6,7]. In the presence of Hh, however, PTCH is inhibited, which allows the SUFU-GLI complex to be recruited into the primary cilium [8]. SUFU dissociates from GLI allowing the translocation of GLI to the nucleus where it acts as a transcriptional activator [7]. In vertebrates there are three GLI proteins, with GLI1 serving solely as an activator as *Gli1* is a Hh target gene, present only after activation of the pathway [6]. GLI2 and GLI3 are either transcriptional activators or repressors, with GLI3 acting primarily as a repressor [6]. Thus, while Hh target gene transcription is tightly regulated through GLI, other proteins regulate GLI’s. One of these is SUFU, a regulator of GLI processing that is key to Hh pathway activation. Through the reduced processing and stability of the GLI3 repressor [9], *Sufu* loss of function alleles cause constitutive expression of Hh target genes in the absence of a Hh ligand [10–12]. Loss of *Sufu* in humans causes medulloblastoma tumor growth [13,14] and Nevoid Basal Cell Carcinoma Syndrome [14,15], while mouse embryos containing a targeted deletion of *Sufu* are embryonic lethal at E9.5 due to neural tube closure defects [11]. Furthermore, the loss of *Sufu* function in the mouse retina restricts retinal neuron differentiation [12] and in the cerebellum results in developmental delays in neuron differentiation [16] and cerebellum mispatterning [17].

In the present study an *in vitro* system utilizing the mouse P19 embryonal carcinoma cell line was employed to study when the Hedgehog pathway is activated, its requirement during differentiation and the effects of *Sufu* loss of function on neural cell fate. The cell line is readily differentiated into neural lineages in the presence of retinoic acid (RA) [18]. RA causes P19 cells to develop functional neurons and supporting astrocytes [19], and although this is well-known, few studies have investigated the role of Hh signaling in this system [20,21]. Of these, one study showed the overexpression of a constitutively active *Gli2* promotes neuronal differentiation [20], but another revealed the knockdown of *Gli2* causes the same phenotype [21]. Thus, these and many unanswered questions remain, and though Hh signaling is precisely controlled *in vivo*, how SUFU is involved in determining neuron and astrocyte cell fate decisions is not known. To address this shortfall, this study aims to better understand the essential role of SUFU in neuroectodermal differentiation. Using the P19 cell model and timeline investigations, we show when Hh signaling is required during RA-induced neural differentiation, and through genetic ablation of *Sufu*, demonstrate that there is an essential role for SUFU in astrocyte determination.

## 2 Materials & Methods

### 2.1 Cell culture & Differentiation

P19 embryonal carcinoma cells (a gift from Dr. Lisa Hoffman, Lawson Health Research Institute, London, ON) and *Sufu* knockout (*Sufu*^*-/-*^, described below) P19 cells were cultured on adherent tissue culture plates in DMEM containing 5% fetal bovine serum and 1% pen/strep at 37°C and 5% CO_2_. Cells were passaged every 4 days or at 70% confluency, whichever occurred first. To induce neural differentiation, approximately 1.05×10^6^ cells were cultured on bacterial grade Petri dishes for 4 days in 0.5 μM RA to form embryoid bodies (EB)[19]. Next, EBs were resuspended and plated on adherent tissue culture dishes in the presence of 0.5 μM RA, for a total of 10, 14 or 17 days. Untreated controls were grown as described, but without RA. For Smoothened Agonist (SAG; EMD Millipore) and Cyclopamine (Cyc; EMD Millipore) studies, cells were cultured as above but in the presence of 0.5 μM RA plus 10 nM SAG or 10 μM Cyc, or 10 nM SAG alone during EB formation; then replated as described above in the presence of 0.5 μM RA or were left untreated for the SAG-only treatment.

### 2.2 Cas9 plasmid preparation

The pSpCas9(BB)-2A-Puro (PX459) V2.0 (Addgene plasmid: 62988) was used, where sgRNAs for *Sufu* (Supplementary Table 1) were cloned into the PX459 vector [22]. Briefly, sgRNAs were amplified and phosphorylated, the vector digested using the BbsI restriction enzyme, and vector and sgRNA were incubated at room temperature for 1.5 h for ligation. Ligated plasmids were transformed into competent bacteria, colonies were selected, and isolated plasmids were sequenced at the London Regional Genomics Centre (Robarts Research Institute, London, ON) using the U6 primer (Supplementary Table 1).

### 2.3 Knockout lines

PX459-sgRNA plasmid (2μg) was transfected using 10 μL of Lipofectamine 2000 (Invitrogen). After 4 h, media was changed, and cells were grown to 70% confluency for clonal selection. Selection media containing 1 μg/mL puromycin was replaced every 24 h for 4 days, and then cells were grown in complete media without puromycin until ready to be passaged. Knockout genotypes were determined by collecting genomic DNA (Qiagen DNeasy® Blood & Tissue kit, 69504), and performing standard PCR (DreamTaq Master Mix (2X), Thermo Scientific) with *Sufu*-specific primers (Supplementary Table 1). Amplicons were sequenced at the London Regional Genomics Centre and compared to wildtype sequences using pair-wise sequence alignment (Geneious 2021™). KO line #2 sequencing was examined using TIDE analysis (http://shinyapps.datacurators.nl/tide/) [23] to further investigate the nature of the allele. TIDE analysis was unable to interrogate KO line #1 as the length of deletion was outside of the tool parameters.

### 2.4 Overexpression lines

*Sufu* KO #2 was transfected with 2 μg of either 1436 pcDNA3 Flag HA (Addgene plasmid: 10792), a pcDNA Flag HA vector with the *Sufu* open reading frame cloned from pcDNA Su(fu) (CT #116) (Addgene plasmid: 13857)[24] using BamHI (New England Biolabs) and XhoI (New England Biolabs) restriction enzymes, or hGli3 flag3x (Addgene plasmid: 84921) using Lipofecatime 2000 (Invitrogen). Cells were incubated for 4 hours in the presence of plasmid, media was then changed, and cells were grown to 70% confluency. Transfected cells were treated with 500 μg/mL neomycin (G418) selection media every 48 hours for 2 weeks.

### 2.5 Real-time reverse transcriptase PCR

To determine relative mRNA expression levels total RNA was collected from RA-treated cells at various time points using QiaShredder (Qiagen) and RNeasy (Qiagen) kits. cDNA was made using a High-Capacity cDNA Reverse Transcription Kit (Thermo Fisher Scientific) and amplification was performed using gene-specific primers (Supplementary Table 1). RT-qPCR was done using 500 nM of each reverse and forward primers, 10 μL of SensiFAST SYBR No-ROX Mix (Bioline), and 1 μL of cDNA template. Samples were analyzed using a CFX Connect Real-Time PCR Detection System (Bio-Rad) using the comparative cycle threshold method (2^-ΔΔCt^). Relative expression was normalized to *L14* and untreated or wildtype untreated controls to determine fold change expression.

### 2.6 Immunoblot Analysis

Cells were lysed in 2% SDS buffer containing 62.5 mM Tris-HCL pH 6.8, 10% glycerol, 5% mercapto-2-ethanol and 1X Halt Protease Inhibitor Cocktail (Thermo Scientific; 1:200). Proteins were separated using SDS-PAGE, then transferred to PVDF membranes (Bio-Rad, 1620177). Membranes were probed with primary antibodies to β-III-tubulin (1:1000; Cell Signaling Technology), GLI3 (1:1000; R&D Systems), SUFU (1:1000; abcam), GFAP (1:1000; Invitrogen) and β-actin (1:10,000; Santa Cruz Biotechnology) and then with the appropriate HRP-conjugated secondary antibodies (1:10,000, Sigma). Signals were detected using the Immobilon® Classico Western HRP Substrate (Millipore) and imaged using a Chemi Doc Touch System (Bio-Rad).

### 2.7 Immunofluorescence

Cells were differentiated, as stated previously, on coverslips coated with poly-L-lysine hydrobromide (Sigma), fixed with 4% paraformaldehyde for 10 min and permeabilized in 0.2% Triton-X-100 for 10 min at room temperature. Coverslips were incubated for 30 min in 1% Bovine Serum Albumin, 22.52 mg/mL glycine and 0.1% Tween-20, then overnight in a humidity chamber with β-III-tubulin (TUJ1, 1:400; Cell Signaling Technology) and GFAP (1:500; Invitrogen). Incubation in secondary antibodies (1:400), Alexa Fluor™ 660 goat anti-mouse IgG (Invitrogen) and Goat anti-Rabbit IgG Alexa Fluor Plus 488 (Invitrogen) was done in the dark for 1 h at room temperature. Coverslips were mounted onto microscope slides using Slowfade™ Gold antifade reagent with DAPI (Invitrogen) and cells were examined using a Zeiss AxioImager Z1 Compound Fluorescence Microscope at the Integrated Microscopy Facility, Biotron (Western University, London, ON). Images were analyzed using ImageJ.

### 2.8 Flow Cytometry

Cells, differentiated as stated previously, were washed with ice cold PBS without magnesium or chloride (PBS(-/-)) (Thermo Fisher) and dissociated with 0.25% Trypsin. Trypsin was deactivated with DMEM/F12 (Thermo Fisher) containing 10% ES-grade FBS (Thermo Fisher), and cells were centrifuged at 300 g for 5 minutes and reconstituted in PBS(-/-). Cells were counted using a trypan blue assay, separated to approximately 1.0×10^6^ cells per tube, and washed in Flow Cytometry Staining Buffer (10% FBS in PBS(-/-)). Cells were fixed using 4% formaldehyde in PBS(-/-) for 10 min at room temperature, washed with ice cold PBS(-/-) and permeabilized with 0.2% Triton-X-100 in PBS(-/-) for 10 min. Cells were incubated with or without 1 μL of GFAP Monoclonal Antibody (GA5), Alexa Fluor 488, eBioscience™ (Invitrogen) per million cells for 1 h at room temperature, and then washed with PBS (-/-), strained using a 40 μm strainer (Falcon™) and analyzed using a BD FACSCanto II flow cytometer at the London Regional Flow Cytometry Facility (Robarts Research Institute). Data were analyzed using FlowJo (BD).

## 3 Results

### 3.1 Retinoic acid and cell aggregation induces P19 neural differentiation

To induce neural differentiation cells were treated for 4 days in 0.5 μM RA to form EB, which were then plated for a total of 10 days to form neurons or 17 days to form astrocytes (Figure 1A). Differentiation was confirmed with RT-qPCR of *NeuroG1* increasing and subsequently decreasing expression (Figure 1B), and *NeuroD1* and *Ascl1* increasing expression throughout RA treatment (Figure 1C and D, respectively). As expected [25], the neuronal marker, β-III-tubulin was detected following RA treatment at 10 days (Figure 1E) and coinciding with prominent signals for Glial Fibrillary Acid Protein (GFAP), an astrocyte marker, that increased by day 17 (Figure 1E). Neither marker was prominent in cells undergoing spontaneous differentiation in the absence of RA (Figure 1E), and together confirmed that neurons and astrocytes differentiated by RA.

**Figure 1.**
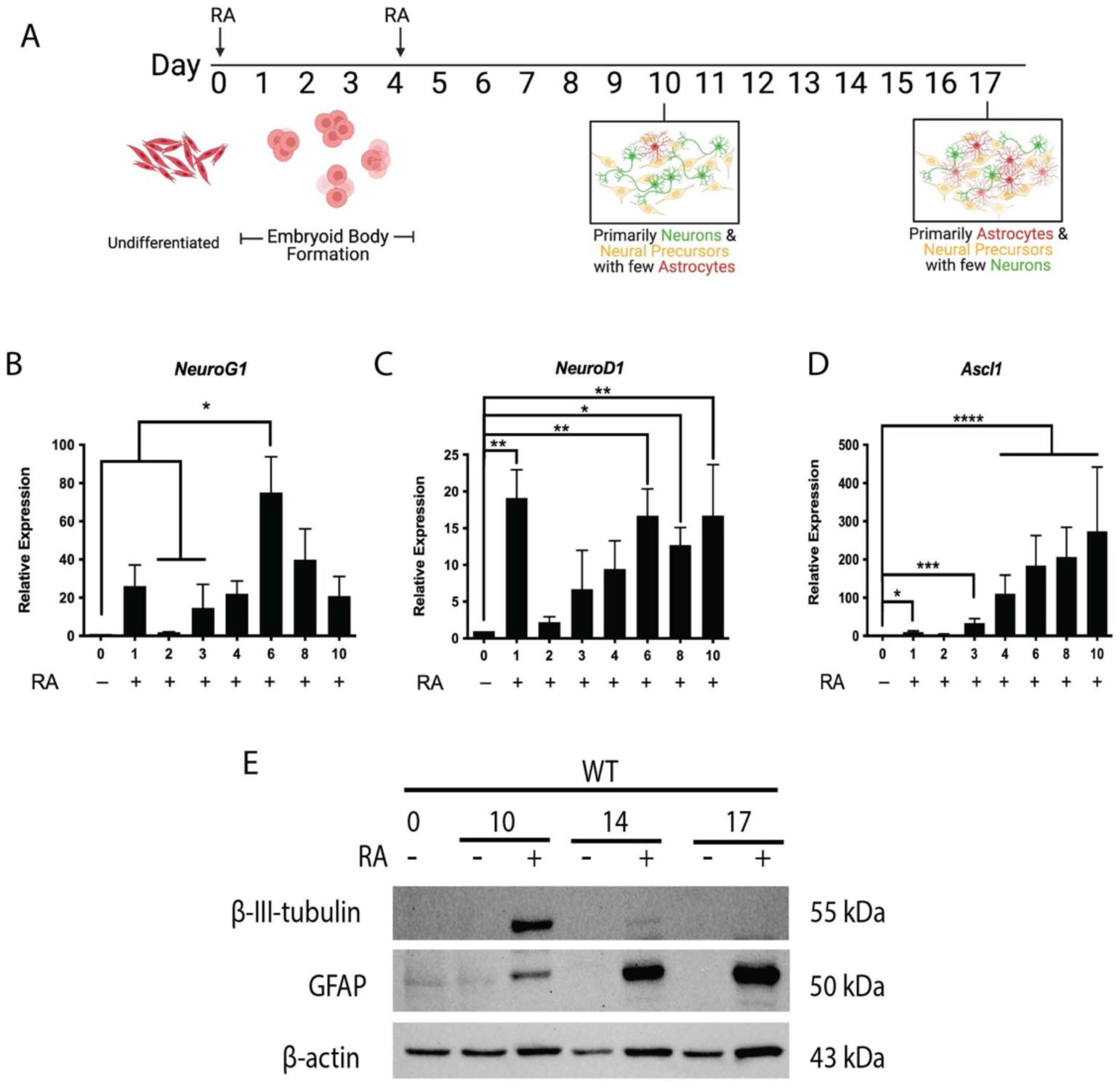
Retinoic acid is required for P19 cell neural differentiation. **(A)** Schematic of *in vitro* differentiation of P19 cells toward neural lineages. RT-qPCR of **(B)** *NeuroG1*, **(C)** *NeuroD1*, and **(D)** *Ascl1* expression on days 0-10 of RA-induced differentiation. **(E)** Immunoblot of β-III-tubulin and GFAP on days 0-17 of RA-induced differentiation. N=3. Bars represent mean values ± s.e.m. *\*P<0*.*05, **P<0*.*01, ***P<0*.*001, ****P<0*.*0001*. Panel A was created using BioRender.com.

### 3.2 Hh signaling is activated early in neural differentiation

Since reports note that Shh signaling is vital in vertebrate neural differentiation [1–3], RT-qPCR was used to explore Hh component expression throughout RA treatment of P19 cells. Hh ligands *Dhh, Ihh*, and *Shh* (Figure 2A-C) were significantly upregulated after 1 day of RA treatment, as was *Gli1* (Figure 2D). Expression of *Ptch1, Ptch2, Smo, Gli2, Gli3* (Supplemental Figure 1), and *Sufu* (Supplemental Figure 3A) were also confirmed. Since GLI3 can serve as an activator or repressor [6] the ratio between GLI3 full-length (GLI3FL) and GLI3 repressor (GLI3R) was examined and results showed an increase in the ratio in untreated cells and RA-treated cells after 1 day in culture (Figure 2E and F). In RA-treated cells GLI3R subsequently increased in abundance (Figure 2E). The increase in GLI3FL to GLI3R in untreated cells was unexpected, as these cells undergoing spontaneous differentiation do not differentiate into neurons or astrocytes (Figure 1E). Together, these results suggest that the EB formation is key to the activation of Hh signaling. At time points associated with astrocyte differentiation (days 10-17), the GLI3FL to GLI3R ratio in RA-treated cells was not significantly different from day 0 controls (Figure 2G). Given these data, it would appear the Hh pathway is activated early in differentiation of these cells, due to the formation of EB, leading to GLI3 activation. However, this activation is reduced for the remainder of RA-induced differentiation.

**Figure 2.**
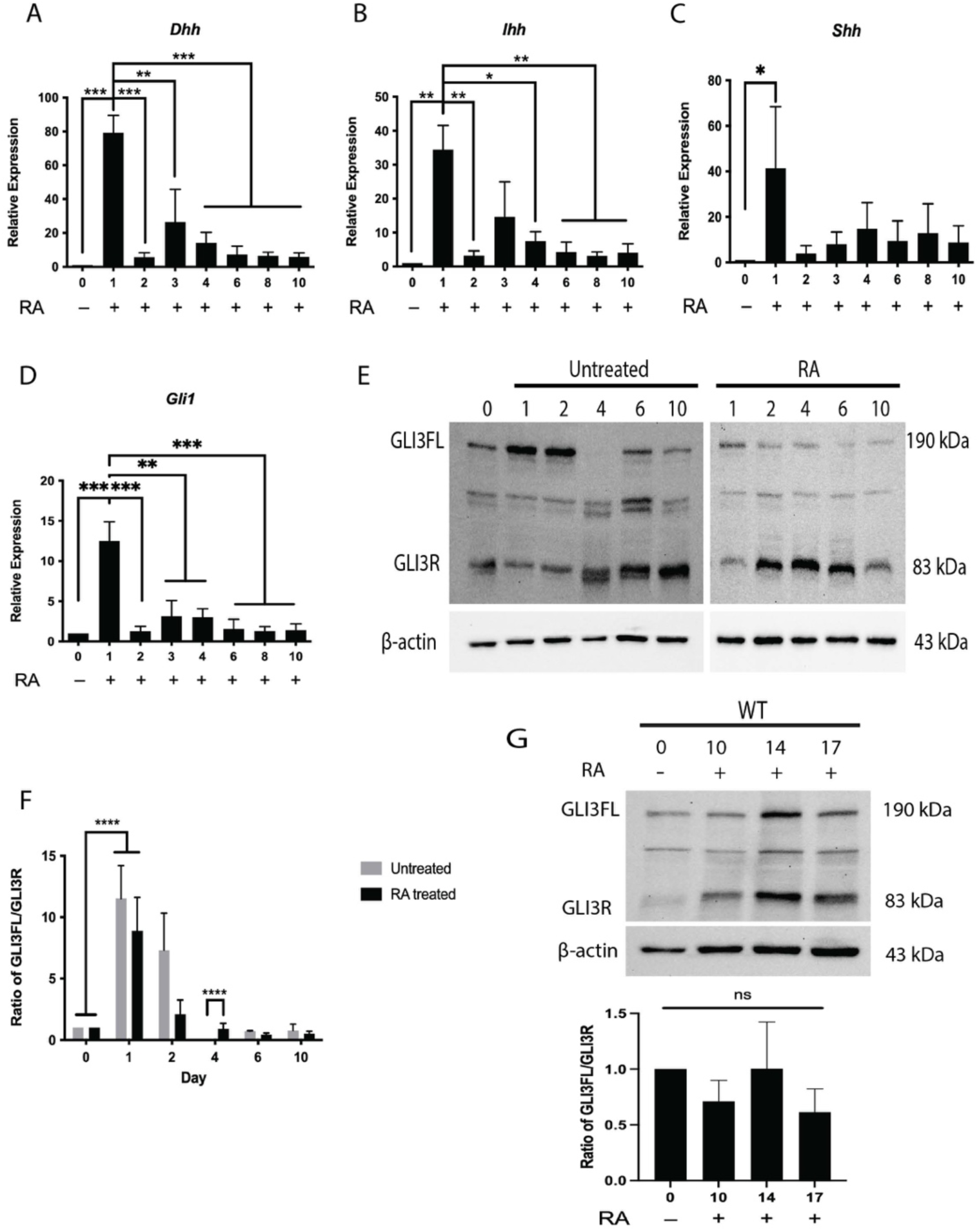
Hh signaling is activated early in RA-induced differentiation. RT-qPCR showing expression of Hh ligands **(A)** *Dhh*, **(B)** *Ihh*, and **(C)** *Shh*, and Hh target gene **(D)** *Gli1* on days 0-10 of RA-induced differentiation. **(E)** Immunoblot of GLI3 full length (GLI3FL) and GLI3 repressor (GLI3R) on days 0-10 of untreated or RA-induced cells. **(F)** Densitometric analysis of GLI3FL to GLI3R ratio of previous immunoblot. (**G)** Upper panel: Immunoblot of GLI3FL and GLI3 repressor on days 0-17 of RA-induced cells, lower panel: densitometric analysis. N=3. Bars represent mean values ± s.e.m. P-values were determined by One-way or Two-way ANOVA, *\*P<0*.*05, **P<0*.*01, ***P<0*.*001, ****P<0*.*0001*.

### 3.3 Early Hh signaling is required but not sufficient for neural differentiation

To first address if activation of Hh signaling would affect differentiation, cells were treated with 0.5 μM RA (RA), 0.5 μM RA plus 10 nM SAG (RA + SAG), 10 nM SAG alone (no RA treatment on day 4) (SAG) or 10 nM SAG on day 0 with subsequent 0.5 μM RA treatment on day 4 (SAG + RA). Immunoblots showed β-III-tubulin in RA, RA + SAG and SAG + RA treatments but not in SAG treatment alone on day 10 (Figure 3A). Densitometric analysis revealed an increase in β-III-tubulin intensity in the RA + SAG treatment compared to RA alone, but no difference between RA alone and the SAG + RA treatment (Figure 3B). Experiments allowing cell aggregation for 4 days in the absence of RA, then subsequent 0.5 μM RA treatment at day 4 (Untr + RA), also showed the β-III-tubulin signal at day 10, however it was significantly less than the RA treatment alone (Supplemental Figure 2A and B). The presence of a β-III-tubulin on day 17 of the blot shown is not representative of the data as densitometric analysis (Figure 3B) showed levels which were not significantly different in cells at all treatments at this stage. Therefore, SAG over-activation of Hh signaling, in the presence of RA, appeared to enhance the neuron cell fate.

**Figure 3.**
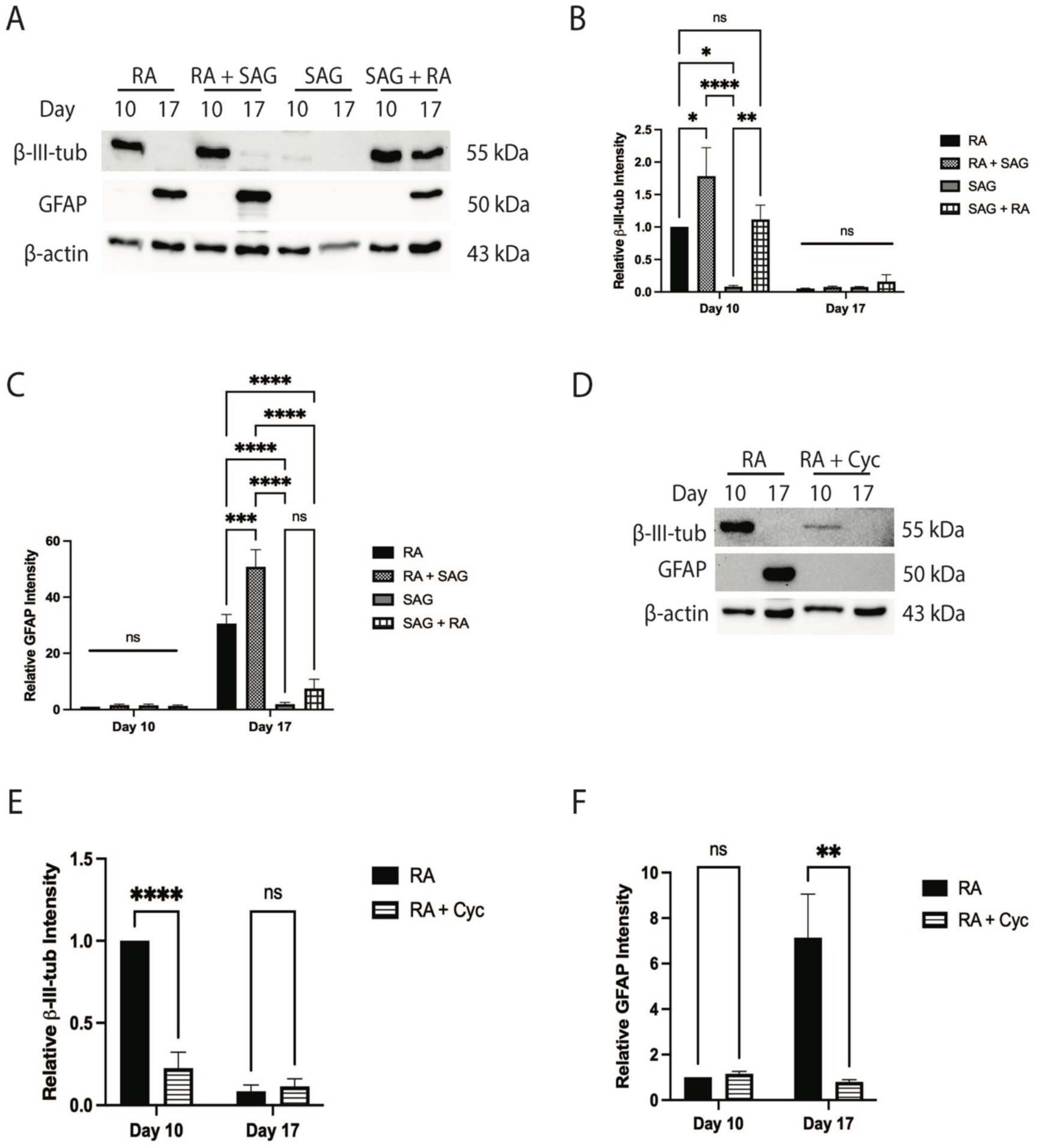
Early Hh pathway activation is required but not sufficient to induce neural lineages. **(A)** Immunoblot of β-III-tubulin and GFAP on days 10 and 17 of cells treated with RA alone, RA plus SAG, SAG alone or SAG plus subsequent RA treatment. **(B)** Densitometry of β-III-tubulin and **(C)** of GFAP from immunoblot in A. **(D)** Immunoblot of β-III-tubulin and GFAP on days 10 and 17 of cells treated with RA or RA plus Cyclopamine (Cyc). **(E)** Densitometry of β-III-tubulin and **(F)** GFAP from immunoblot in D. N=3. Bars represent mean values ± s.e.m. P-values were determined by Two-way ANOVA, *\*P<0*.*05, **P<0*.*01, ***P<0*.*001, ****P<0*.*0001*.

Immunoblot analysis of GFAP showed detection on day 17 in the RA and RA + SAG treatments, but no signals were seen in the SAG alone treatment (Figure 3A). Furthermore, densitometric analysis showed increased GFAP intensity in the RA + SAG treatment compared to RA treatment alone (Figure 3C). A GFAP band was also seen in the SAG + RA treatment (Figure 3A), but as described above, densitometric analysis revealed that this signal was not significantly different from the SAG alone treatment (Figure 3C). Untr + RA cells, also showed this low level of GFAP at day 17 (Supplemental Figure 2A and C), indicating that RA treatment at this later time was sufficient to induce astrocytes, however not to the extent observed with the initial RA treatment. Thus, it appears from immunoblot data that supplementing the initial RA treatment with SAG increases astrocyte differentiation, but activation of Hh signaling alone using SAG was not sufficient to induce astrocyte fate.

Since early Hh pathway activation enhances neural cell fates, would its attenuation affect neural differentiation? Cells were treated with either 0.5 μM RA (RA) or 0.5 μM RA plus 10 μM Cyc on day 0 (RA + Cyc); both with subsequent RA treatments on day 4. Immunoblot analysis of β-III-tubulin on day 10 showed signals in both RA and RA + Cyc treatments (Figure 3D), but densitometry revealed the intensity was significantly reduced with Cyc treatment (Figure 3E). Cyc had dramatic effects on GFAP levels, eliminating the response induced by RA (Figure 3D and F). Experiments with cells treated with RA plus the Cyc vehicle (DMSO) showed no significant change in either marker (Supplemental Figure 2D-F). These results suggest that inhibition of the Hh pathway attenuates neuron and astrocyte cell fates, and thus is required for proper differentiation of these cell types.

#### SUFU levels during neural differentiation

SUFU negatively regulates GLI activity and as reported by Humke et al. [7], promotes GLI3 conversion to a repressor. To determine its role in Hh signaling seen in P19 cells, *Sufu* gene expression was examined and found to increase on day 1, 8 and 10 of RA treatment (Supplemental Figure 3A); however, at the protein level no changes were seen between untreated and RA treated cells, and no changes were detected over time by either treatment (Supplemental Figure 3B and C). Similarly, *Sufu* expression on days 10-17 showed no significant change relative to day 0 (Supplemental Figure 3D); however, SUFU protein intensity decreased at these later stages (Supplemental Figure 3E and F). Since *Sufu* expression remains largely unchanged and only at later stages, coinciding with astrocyte differentiation, do protein levels decrease, it is tempting to speculate that the early activation of the Hh pathway is controlled by the cellular localization or post-translational modification status of SUFU.

#### Sufu knockout activates Hedgehog signaling

Since SUFU acts as an essential regulator of GLI3 [7] and GLI2 [4,7], its loss was expected to induce Hh signaling. To test this, CRISPR-Cas9 was used to target exon 6 of the *Sufu* gene (Figure 4A), resulting in clones with an 81 bp deletion (KO #1) that spans the intron-exon boundary between intron 5 and exon 6 and another with a 1 bp insertion (KO #2). Sequence alignment, TIDE analysis [23] and *in silico* translation of clones revealed a premature stop codon shortly after the mutation (Supplemental Figure 4). Immunoblot analyses using a SUFU antibody targeted to the C-terminus of SUFU, a region reported to be required for GLI binding [4,26], was detected in untreated WT cells, but was not in either mutant clone (Figure 4B). To determine other effects, *Sufu*^*-/-*^ alleles caused a decrease in cell numbers compared to WT cells cultured to 96 h (Figure 4C); however, MTT analysis at the same timepoints showed no change in relative MTT absorbance (Figure 4D). As predicted, KOs showed increased expression of Hh targets *Gli1, Ptch1*, and *Ascl1* in undifferentiated cultures (Figure 4E). Thus, the data shows the loss of SUFU does not greatly affect the proliferation of undifferentiated cells, contrary to that reported in mouse neural stem cells of the dentate gyrus [27], and instead it induced Hh target genes in P19 cells.

**Figure 4.**
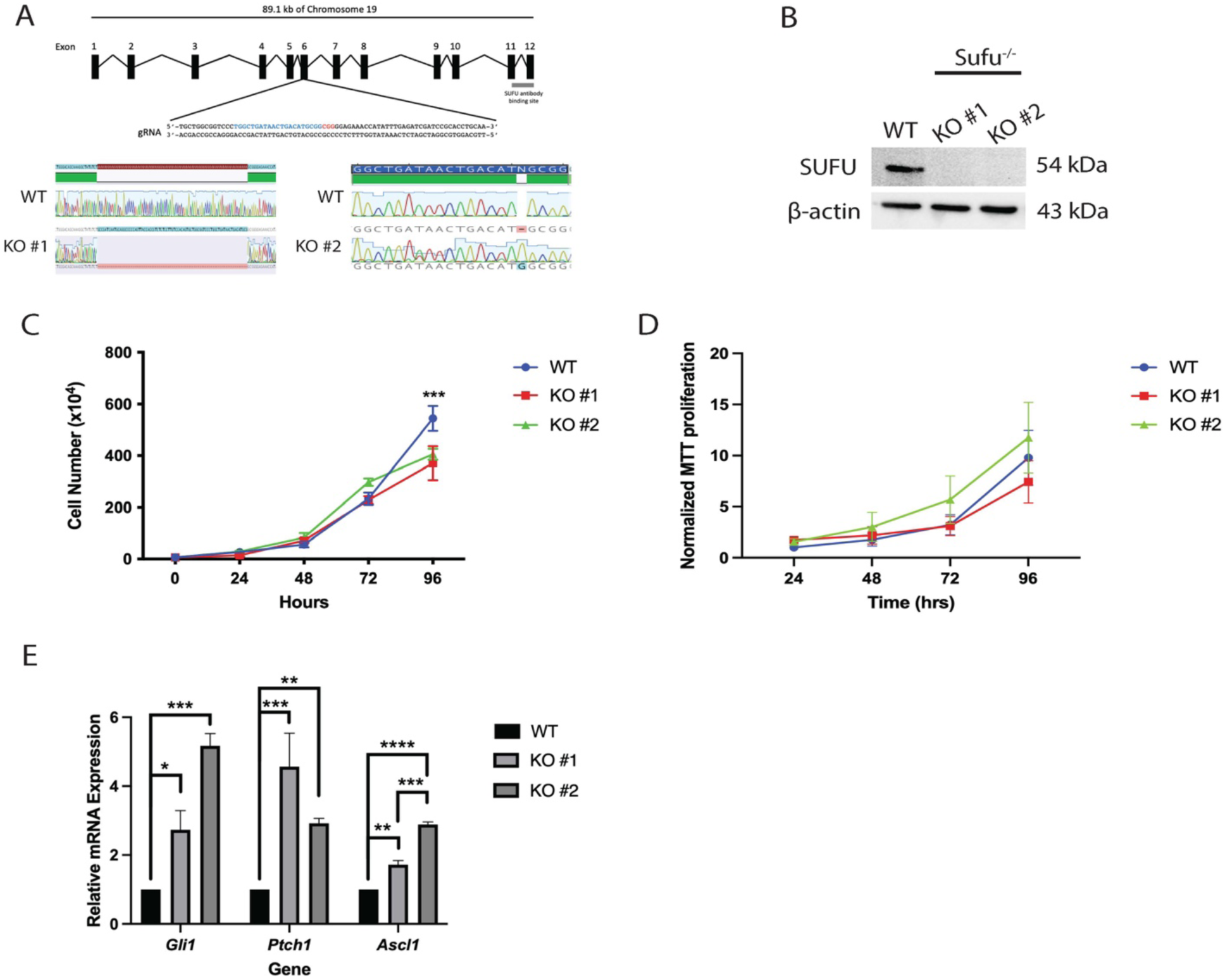
CRISPR-Cas9 knock out of *Sufu* activates Hh signaling in undifferentiated P19 cells. **(A)** Schematic showing gRNA targeting exon 6 of the *Sufu* gene yielding two KO clones. **(B)** Immunoblot of SUFU levels in wildtype (WT) and *Sufu*^*-/-*^ clones. **(C)** Cell count curves of WT and *Sufu*^*-/-*^ clones between 0-96 hours and **(D)** MTT assay normalized to WT cells at 24 hrs. **(E)** Expression of *Gli1, Ptch1* and *Ascl1* in WT and *Sufu*^*-/-*^ clones on day 3 of undifferentiated cultures. N=3. Bars represent mean values ± s.e.m. P-values were determined by either One-way or Two-way ANOVA, *\*P<0*.*05, **P<0*.*01, ***P<0*.*001, ****P<0*.*0001*.

### 3.4 Sufu loss activates Hh signaling through loss of Gli3

RT-qPCR was used to determine if the observed activation of Hh signaling in the KO lines was maintained throughout RA-induced differentiation. *Sufu*^*-/-*^ cells treated with RA maintained higher *Gli1* (Figure 5A and Supplemental Figure 5A) and *Ptch1* (Figure 5B and Supplemental Figure 5B) expression compared to that in WT cells. Since previous work showed the connection between SUFU and GLI3 processing [7], we were particularly interested in *Gli3* expression and protein levels. Results show no significant differences in *Gli3* expression between *Sufu*^*-/-*^ and WT cells in KO#2 (Figure 5C), however, significantly decreased *Gli3* levels were observed at days 10, 14 and 17 in KO #1 (Supplemental Figure 5C). At the protein level, neither the GLI3 repressor nor full-length GLI3 were detected in KO #2 (Figure 5D), and this was consistent with KO #1 having decreased or absent GLI3 levels (Supplemental Figure 5D). Thus, the loss of SUFU, specifically the region of the polypeptide required for GLI3 processing, likely contributes to post-translational perturbations to GLI3, causing its loss and the observed higher expression of Hh target genes.

**Figure 5.**
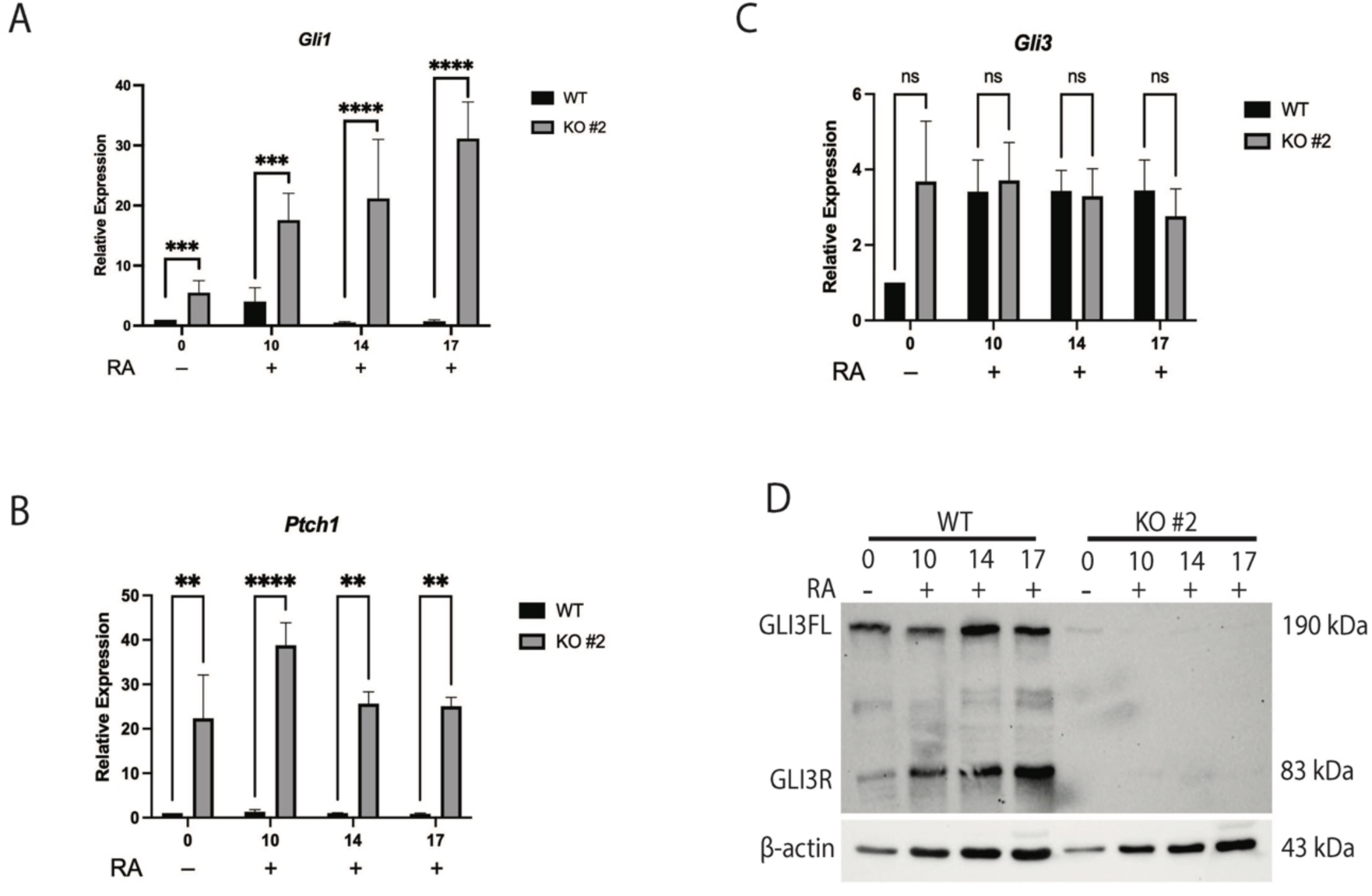
*Sufu*^-/-^ maintains activate Hh signaling and causes the loss of Gli3. Expression of **(A)** *Gli1* and **(B)** *Ptch1* and **(C)** *Gli3* in wildtype (WT) and *Sufu*^*-/-*^ KO #2 cells on days 0-17. **(D)** Immunoblot of GLI3 full length (GLI3FL) and GLI3 repressor (GLI3R) in WT and *Sufu*^*-/-*^ KO #2 cells on days 0-17. N=3. Bars represent mean values ± s.e.m. P-values were determined by Two-way ANOVA, *\*P<0*.*05, **P<0*.*01, ***P<0*.*001, ****P<0*.*0001*.

### 3.5 Sufu knockout delays and decreases the astrocyte cell fate

If the *Sufu* knockout induced Hh target genes what effect would this have on neural differentiation? To determine if KO cells could differentiate in the absence of RA, these cells were aggregated and replated in the absence or presence of RA. Both KO lines treated with RA showed β-III-tubulin signals at day 10 and GFAP signals on days 14 and 17; neither marker was detected in the absence of RA (Figure 6A). To determine if the timing or degree of differentiation was affected by the loss of SUFU, the expression of these markers was compared in each knockout line with that in WT cells at days 10, 14 and 17 in the presence of RA (Figure 6B). Densitometric analyses confirmed β-III-tubulin levels did not change between WT and KO cells (Figure 6C). Significantly, astrocyte differentiation was affected and GFAP signals on immunoblots were absent on day 10 and reduced on day 14 of RA treatment in *Sufu*^*-/-*^ cells compared to WT (Figure 6B and D). The absence of GFAP on day 10 was interpreted cautiously, as in WT cells, we have seen inconsistencies in GFAP levels at this time point. Nevertheless, GFAP levels are consistently high in WT day 14 cells (Figure 1E and Figure 8C) and it appeared removing SUFU delayed astrocyte differentiation when RA was present.

**Figure 6.**
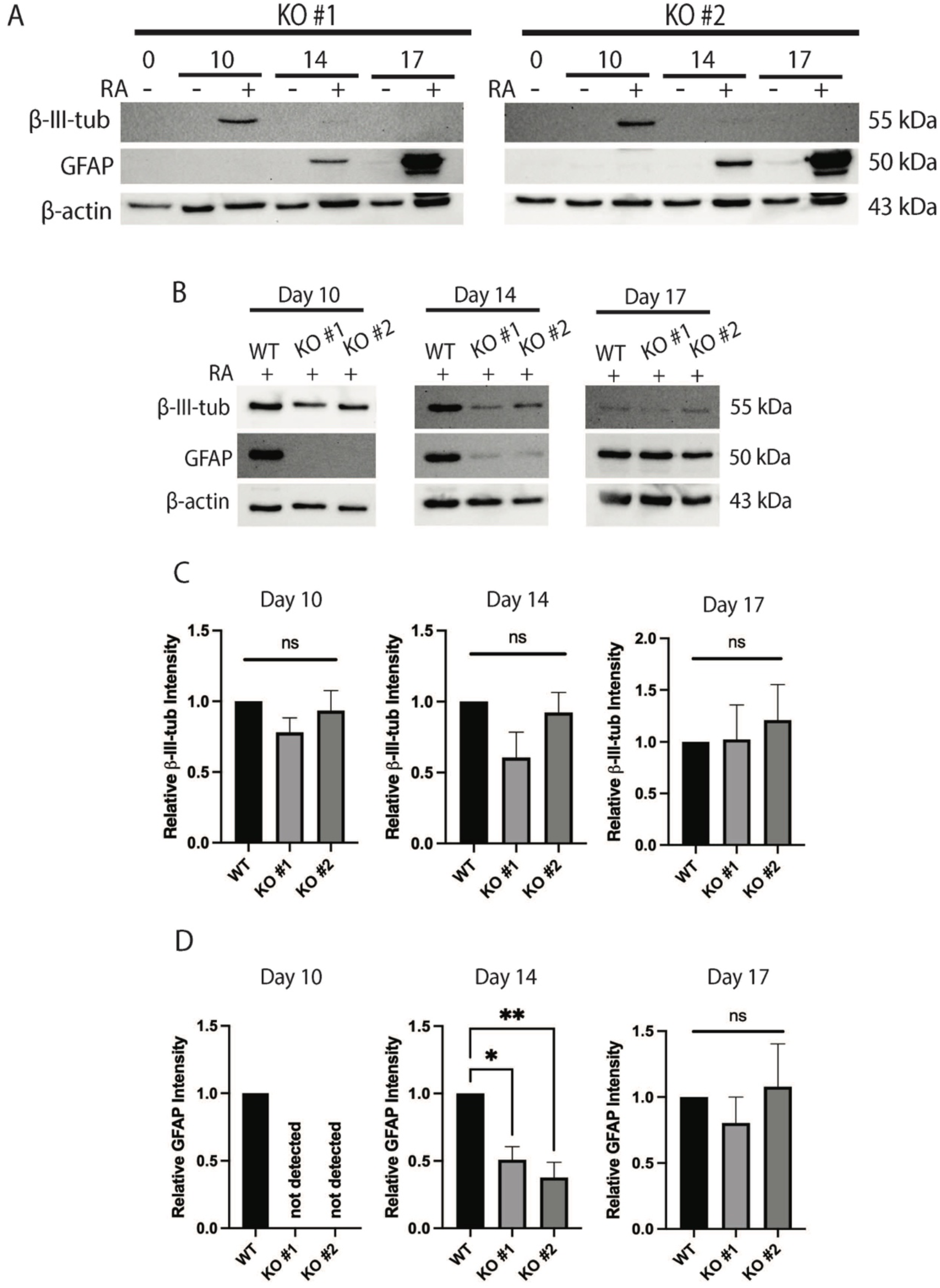
*Sufu*^*-/-*^ delays astrocyte formation without affecting neuron differentiation. **(A)** Immunoblot of ? β-III-tubulin and GFAP on days 0-17 in *Sufu*^*-/-*^ KO clones. **(B)** Immunoblots of β-III-tubulin and GFAP on day 10, day 14 and day 17 in wildtype (WT) and *Sufu*^*-/-*^ clones, and densitometric analyses of immunoblots of B) showing relative intensities of **(C)** β-III-tubulin and **(D)** GFAP. N=3. Bars represent mean values ± s.e.m. P-values were determined by One-way ANOVA, *\*P<0*.*05, **P<0*.*01*.

To confirm the immunoblot data showing astrocyte differentiation was restored in *Sufu*^*-/-*^ cells on day 17 (Figure 6D), an immunofluorescence analysis with antibodies to β-III-tubulin and GFAP were used on WT and *Sufu*^*-/-*^ cells. Although expecting to see no differences in either marker between the two cell lines, this analysis revealed a reduction in GFAP-positive staining in *Sufu*^-/-^ cells (Figure 7A). However, since these cells are highly heterogenous and exist in areas of high-density, flow cytometry was used and results confirmed this reduction showing *Sufu*^*-/-*^ cells (∼6-25%) had significantly less GFAP-positive staining compared to WT cells (∼60-70%; Figure 7B and C). It should be noted that although the immunoblot data at day 17 was inconsistent with the immunofluorescence and flow analysis, the flow cytometry had identified high GFAP levels (Figure 7B) in some *Sufu*^*-/-*^ cells, even though overall there were fewer GFAP positive cells present in the KO line (Figure 7C). Nevertheless, these data support the idea that the loss of *Sufu* does not alter neuronal differentiation, but it does delay and decrease astrocyte differentiation.

**Figure 7.**
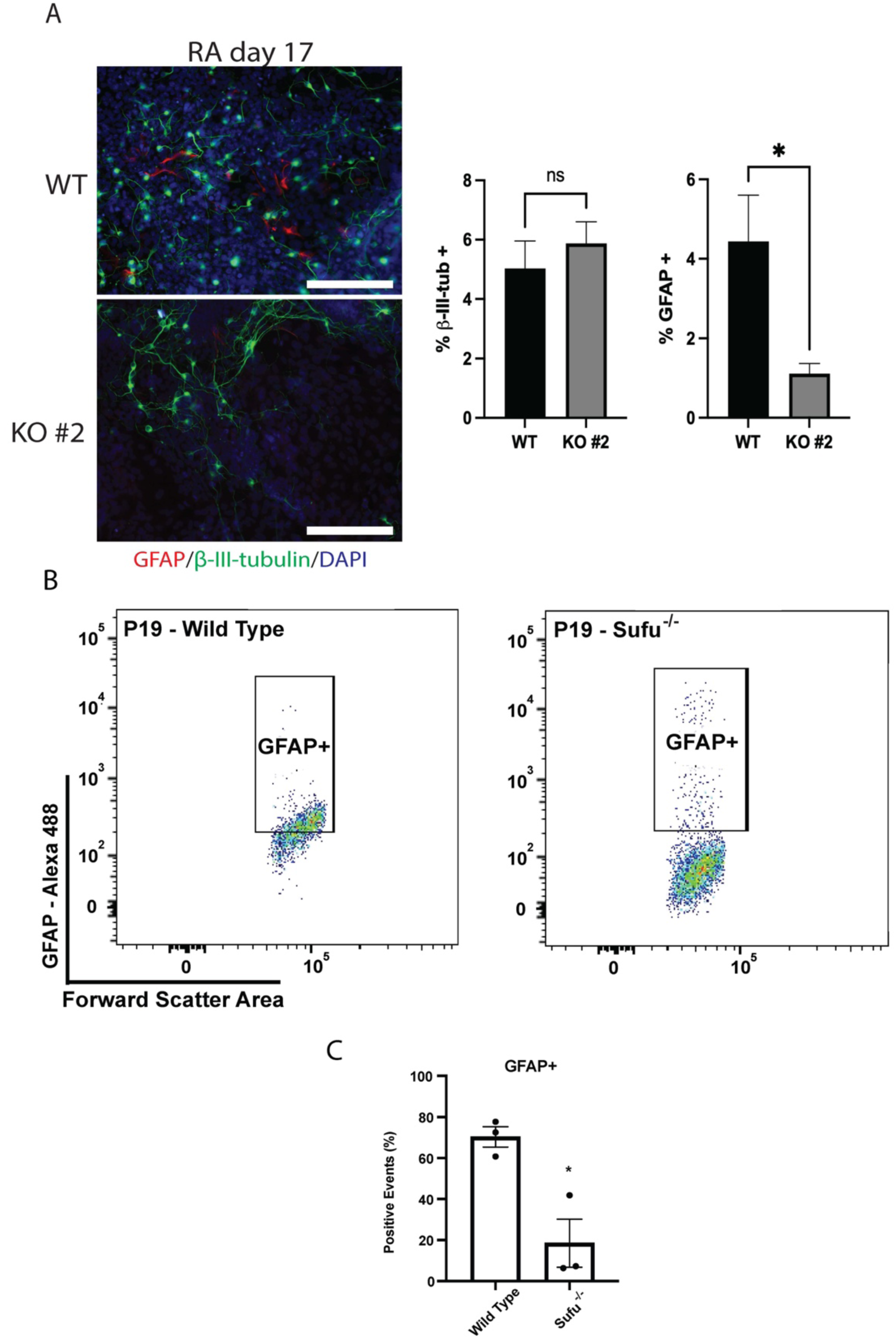
*Sufu*^*-/-*^ decreases astrocyte differentiation. **(A)** GFAP (red), β-III-tubulin (green) and nuclear DAPI (blue) fluorescence in wildtype (WT) and *Sufu*^*-/-*^ KO clone #2 cells on day 17. Scale bar = 200 μm. **(B)** Flow cytometry of GFAP-positive WT and *Sufu*^*-/-*^ KO #2 cells on day 17 and quantified in **C**. N=3. P-values were determined by Student’s t-test, *\*P<0*.*05*.

### 3.6 Sufu or Gli3 overexpression partially rescues astrocyte phenotype

If the loss of *Sufu* resulted in delayed and decreased astrocyte differentiation (Figure 6 and Figure 7) and this was likely due to the loss of GLI3 (Figure 5D and Supplemental Figure 5D), then overexpression of a CRISPR-insensitive chicken *Sufu* (pSUFU) or human *GLI3* (pGli3) in *Sufu* knockout cells should rescue the phenotype and astrocyte levels should increase in these mouse cells. Overexpression was confirmed by RT-qPCR of chicken *Sufu* and immunoblotting of human GLI3 (Supplemental Figure 6). To determine the effect this overexpression had on astrocyte differentiation, GFAP levels were explored following pSUFU or pGli3 stable transfection in knockout cells. Results showed no GFAP detection on day 10, but there was an increase by day 14 of RA treatment, albeit not to the levels observed in the WT cells (Figure 8A and B). GFAP levels were maintained at day 17 in the human pGli3 stably transfected cells, but chicken pSUFU stably transfected cells showed significantly reduced levels compared to the WT or other transfected *Sufu*^*-/-*^ cells (Figure 8A and B). Despite these minor differences, these results show that chicken *Sufu* and human *GLI3* overexpression partially rescue astrocyte differentiation and highlights not only the essential role of *Gli3* in neural differentiation, but a major role for SUFU in regulating GLI3.

**Figure 8.**
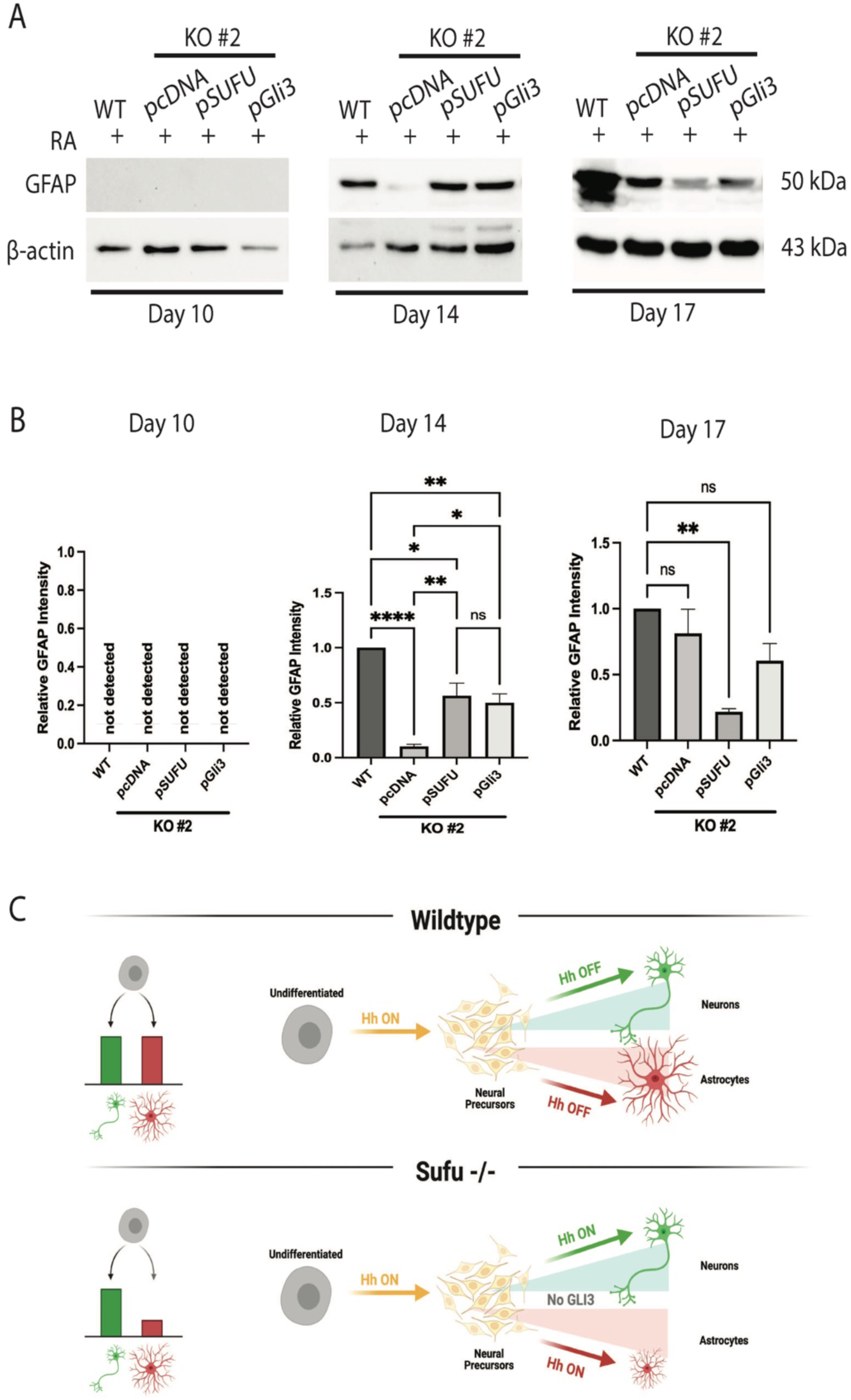
*Sufu* or *Gli3* overexpression partially rescues *Sufu*^*-/-*^ phenotype. **(A)** Immunoblot of GFAP on days 10, 14 and 17 in WT and *Sufu*^*-/-*^ KO clone #2 cells stably transfected with pcDNA empty vector (pcDNA), pcDNA-chicken Sufu (pSUFU), or pcDNA-human Gli3 (pGli3) vectors and treated with RA. Densitometric analysis of **(B)** GFAP from the immunoblot in A. N=3. Bars represent mean values ± s.e.m. P-values were determined by One-way ANOVA, *\*P<0*.*05, **P<0*.*01, ****P<0*.*0001*. **(C)** Summary comparing WT and *Sufu*^*-/-*^ cell neural differentiation.

## 4 Discussion

Hh signaling is involved in neural tube closure and neural differentiation *in vivo* [1,3] where its activation results in transcription of target genes through GLIs [3,7]. Negative regulators of the pathway include Patched and SUFU [3], the loss of which results in medulloblastoma and Nevoid Basal Cell Carcinoma Syndrome [13–15,28]. Using the mouse P19 cell model, and summarized in Figure 8C, we report the early activation of the Hh pathway is required for neural differentiation, but its activation alone is not sufficient to induce neural cell fates. Furthermore, SUFU was found to be essential for *in vitro* neural differentiation as its loss activated Hh signaling through the loss of Gli3, causing perturbed timing and proportion of astrocyte differentiation. Furthermore, human *Gli3* overexpression was sufficient to rescue GFAP, the astrocyte marker, in SUFU deficient mouse cells, which indicates it being downstream or parallel to the negative regulation imparted by SUFU.

RA-induced neural differentiation of P19 cells is well established [18,19,25,29], and few studies have explored Hh signaling in the model [20,21,30]. RA likely acts in these cells through the transcription factor FOXA1, which promotes the early expression of *Shh* [30], but the precise timing of pathway activation throughout differentiation has not been heretofore reported. We have shown that EB formation with and without RA triggers early Hh pathway activation (Figure 2). Although the mechanism of activation in untreated cells remains unclear, these cells do not spontaneously differentiate towards neural lineages in the absence of RA (Figure 1E). Focusing on RA-treated cells we demonstrated that RA induced *Shh* expression, but *Ihh* and *Dhh* were also expressed (Figure 2). IHH and DHH, like SHH, can activate the pathway as evident in long bone differentiation [31,32] and gonad development [33–35]. Similar results showing transcriptional activation of all Hh ligands was reported with human embryonic stem cells (hESCs) induced to form neural tissues [36]. Thus, it is likely that SHH is responsible, but since transcripts encoding all ligands are induced by RA it remains unclear which isoform triggers pathway activation in the P19 model. Although we, like others, have demonstrated the presence of Hh signaling in P19 differentiation, other signaling pathways including those linked to Wnt, Fgf and Yap are also involved [21,37–39]. Considering our study, these other pathways require further investigation to determine if they too are temporally regulated to facilitate differentiation.

Utilization of RA to induce neural lineages *in vitro* goes beyond the P19 model, as RA is used for directed neural differentiation of human induced pluripotent stem cells and mouse embryonic stem cells (mESCs) [40,41]. Many neural differentiation protocols also supplement RA with exogenous SHH or SAG in order to enhance the differentiation of neurons [41–43] and glial cell progenitors [44]. Like these previous results, co-treating P19 cells with RA and SAG enhanced the differentiation of both neurons and astrocytes (Figure 3), although treating with SAG alone was not sufficient to induce differentiation. We also investigated the effect of inhibiting the Hh pathway using Cyc, where Cyc treatment inhibited neuron and astrocyte differentiation (Figure 3), a result consistent with previous work showing Cyc co-treatment with RA decreased expression of neural-related transcription factors [43]. Thus, the early activation of Hh signaling in the differentiation of neural lineages is a requirement, and we next sought to understand the importance of later-stage inhibition of the Hh pathway by SUFU.

SUFU regulation of Hh signaling involves the ubiquitin-proteasome pathway following Hh pathway activation [45]. We showed Hh ligand expression is induced within 1 day of RA treatment (Figure 2) and given previous reports on SUFU regulation, an early decrease in SUFU levels was expected. Our results, however, showed no significant change in SUFU levels at early stages, although there was a relative decrease later during astrocyte differentiation (Supplemental Figure 3). The mechanism of SUFU depletion was not determined, nor were SUFU post-translational modifications examined, but it should be noted that differential phosphorylation and ubiquitination can both promote or inhibit SUFU stability and/or GLI interaction [46–49]. CRISPR-Cas9, used to deplete SUFU from P19 cells, show the loss of SUFU induces the ectopic transcription of Hh target genes (Figure 4 and Figure 5) and while supported by studies using mice, mESCs and hESCs [15,36,50–52], this is in contrast with a study showing the reduction in expression in adult neural stem cells [27]. Interestingly, SUFU-deficient P19 cells did not differentiate in the absence of RA (Figure 6A) despite a previous report that *Gli2* overexpression in P19 cells was sufficient to induce neuron differentiation under similar conditions [20]. That another showed the knockdown of *Gli2* causes the same phenotype [21], would indicate that other GLIs are involved, specifically GLI3.

The connection between aberrant astrocyte differentiation and SUFU deficiency came from examining the transcriptional activation of Hh target genes (Figure 4 and Figure 5). *Gli1* and *Ptch1* expression was higher when *Sufu* was knocked out, but not *Gli3* (Figure 5). Previous reports with mouse show that SUFU is required for proper GLI3 processing [53] and its absence leads to reduced or absent GLI3 protein [9,50], which also occurs with conditional *Sufu* knockouts in the mouse brain [16,17,54,55]. We reported similar results with GLI3 in P19 cells and showed that it affects neural differentiation. In support, conditional knockouts of *Sufu* at various stages of brain development also results in decreased neuron differentiation and disorganized glial cell arrangements [16,54]. Discrepancies, however, do exist even within Hh signaling in P19 cells [20,21], and although these differences can be explained, our work has shown the knockout of *Sufu* was accompanied by the consistent activation of Hh target genes, like that seen in mouse retinal progenitor cells lacking SUFU [12]. This activation did not result in neuron differentiation without RA (Figure 6) and is supported by Hoelzl et al. [51], using mESCs depleted of SUFU. Our results did show a delay and decrease in astrocyte differentiation with SUFU loss (Figure 6 and Figure 7), which may explain the disorganization of glial cells observed in conditional *Sufu* knockouts described in mouse embryos [16,54]. Overall, our research shows that Hh signaling is induced by RA, but the pathway must remain in the *off* state for proper astrocyte differentiation. Given that glial cell populations are composed of astrocytes, oligodendrocytes and microglia, further research must be done to explore the effect of SUFU loss on these other lineages. In addition to the delay in astrocyte formation (Figure 6 and 7), there was a loss of GLI3 in these cells (Figure 5) and we have provided evidence that ectopic expression of GLI3 was sufficient to rescue the timing of glial cell differentiation (Figure 8). That *GLI3* was able to rescue the *Sufu* knockout phenotype is significant and supported by previous research showing *Gli3* repressor overexpression in the SUFU-deficient mouse cerebellum was sufficient to rescue morphology and increase GFAP levels [17]. Together, our results clearly show that SUFU imparts its negative regulation on the Hh pathway by acting on Gli3 and this transcription factor is required later during the specification of neural cells to astrocytes.

## 5 Conflict of Interest

*The authors declare that the research was conducted in the absence of any commercial or financial relationships that could be construed as a potential conflict of interest*.

## 6 Author Contributions

Conceptualization, D.M.S. and G.M.K.; methodology, D.M.S.; flow cytometry analysis, J.G.D.; formal analysis, G.M.K. and D.M.S.; writing, first draft—D.M.S.; writing—review and editing, D.M.S. and G.M.K.; supervision, G.M.K.; funding acquisition, G.M.K. All authors have read and agreed to the published version of the manuscript.

## 7 Funding

This research was funded by the Natural Sciences and Engineering Research Council (NSERC) of Canada, Discovery Grant R2615A02 to G.M.K. D.M.S. acknowledges support from the School of Graduate and Postdoctoral Studies, University of Western Ontario; the Collaborative Graduate Specialization in Developmental Biology, University of Western Ontario; the Children’s Health Research Institute; and NSERC for a PGS D scholarship.

## 8 Acknowledgments

We would like to thank members past and present of the Kelly lab for helpful discussions.

## Notes

### Competing Interest Statement

The authors have declared no competing interest.

### Summary of Updates

Figure 8C has been added, and written changes for clarity.

## References

1. Wilson L and M Maden (2005). The mechanisms of dorsoventral patterning in the vertebrate neural tube. Dev Biol 282: 1–13.

2. Dahmane N and A Ruiz (1999). Sonic hedgehog and cerebellum development. Development 3100: 3089–3100.

3. Briscoe J and PP Thérond (2013). The mechanisms of Hedgehog signalling and its roles in development and disease. Nat Rev Mol Cell Biol 14: 418–431.

4. Han Y, Q Shi and J Jiang (2015). Multisite interaction with Sufu regulates Ci/Gli activity through distinct mechanisms in Hh signal transduction. Proc Natl Acad Sci 112: 6383–6388.

5. Tempe D, M Casas, S Karaz, M-F Blanchet-Tournier and J-P Concordet (2006). Multisite Protein Kinase A and Glycogen Synthase Kinase 3 Phosphorylation Leads to Gli3 Ubiquitination by SCF TrCP. Mol Cell Biol 26: 4316–4326.

6. Hui C and S Angers (2011). Gli proteins in development and disease. Annu Rev Cell Dev Biol 27: 513–537.

7. Humke EW, K V. Dorn, L Milenkovic, MP Scott and R Rohatgi (2010). The output of Hedgehog signaling is controlled by the dynamic association between Suppressor of Fused and the Gli proteins. Genes Dev 24: 670–682.

8. Haycraft CJ, B Banizs, Y Aydin-Son, Q Zhang, EJ Michaud and BK Yoder (2005). Gli2 and Gli3 localize to cilia and require the intraflagellar transport protein Polaris for processing and function. PLoS Genet 1: e53.

9. Makino S, O Zhulyn, R Mo, V Puviindran, X Zhang, T Murata, R Fukumura, Y Ishitsuka, H Kotaki, D Matsumaru, S Ishii, CC Hui and Y Gondo (2015). T396I mutation of mouse Sufu reduces the stability and activity of Gli3 repressor. PLoS One 10: 1–15.

10. Svärd J, KH Henricson, M Persson-Lek, B Rozell, M Lauth, Å Bergström, J Ericson, R Toftgård and S Teglund (2006). Genetic elimination of suppressor of fused reveals an essential repressor function in the mammalian hedgehog signaling pathway. Dev Cell 10: 187–197.

11. Cooper AF, KP Yu, M Brueckner, LL Brailey, L Johnson, JM McGrath and AE Bale (2005). Cardiac and CNS defects in a mouse with targeted disruption of suppressor of fused. Development 132: 4407–4417.

12. Cwinn MA, C Mazerolle, B McNeill, R Ringuette, S Thurig, CC Hui and VA Wallace (2011). Suppressor of fused is required to maintain the multipotency of neural progenitor cells in the retina. J Neurosci 31: 5169–5180.

13. Taylor MD, L Liu, C Raffel, C chung Hui, TG Mainprize, X Zhang, R Agatep, S Chiappa, L Gao, A Lowrance, A Hao, AM Goldstein, T Stavrou, SW Scherer, WT Dura, B Wainwright, JA Squire, JT Rutka and D Hogg (2002). Mutations in SUFU predispose to medulloblastoma. Nat Genet 31: 306–310.

14. Smith MJ, C Beetz, SG Williams, SS Bhaskar, JO Sullivan, B Anderson, SB Daly, JE Urquhart, Z Bholah, D Oudit, E Cheesman, A Kelsey, MG Mccabe, WG Newman, DGR Evans, MJ Smith, SG Williams, E Urquhart, Z Bholah and G William (2018). Germline mutations in SUFU cause Gorlin Syndrome – Associated Childhood Medulloblastoma and redefine the risk associated with PTCH1 mutations. J Clin Oncol 32: 4155–4161.

15. Urman NM, A Mirza, SX Atwood, RJ Whitson, KY Sarin, JY Tang and AE Oro (2016). Tumor-derived suppressor of fused mutations reveal hedgehog pathway interactions. PLoS One 11: 1–10.

16. Kim JJ, T Jiwani, S Erwood, J Loree and ND Rosenblum (2018). Suppressor of fused controls cerebellar neuronal differentiation in a manner modulated by GLI3 repressor and Fgf15. Dev Dyn 247: 156–169.

17. Kim JJ, PS Gill, L Rotin, M van Eede, RM Henkelman, C-C Hui and ND Rosenblum (2011). Suppressor of Fused controls mid-hindbrain patterning and cerebellar morphogenesis via GLI3 repressor. J Neurosci 31: 1825–1836.

18. McBurney MW, EMV Jones-Villeneuve, MKS Edwards and PJ Anderson (1982). Control of muscle and neuronal differentiation in a cultured embryonal carcinoma cell line. Nature 299: 165–167.

19. McBurney MW, KR Reuhl, a I Ally, S Nasipuri, JC Bell and J Craig (1988). Differentiation and maturation of embryonal carcinoma-derived neurons in cell culture. J Neurosci 8: 1063–1073.

20. Voronova A, A Fischer, T Ryan, A Al Madhoun and IS Skerjanc (2011). Ascl1/Mash1 is a novel target of Gli2 during Gli2-induced neurogenesis in P19 EC cells. PLoS One 6:.

21. Lin YT, JY Ding, MY Li, TS Yeh, TW Wang and JY Yu (2012). YAP regulates neuronal differentiation through Sonic hedgehog signaling pathway. Exp Cell Res 318: 1877–1888.

22. Ran FA, PD Hsu, J Wright, V Agarwala, DA Scott and F Zhang (2013). Genome engineering using CRISPR-Cas9 system. Nat Protoc 8: 2281–2308.

23. Brinkman EK, T Chen, M Amendola and B Van Steensel (2014). Easy quantitative assessment of genome editing by sequence trace decomposition. Nucleic Acids Res 42: 1–8.

24. Pearse R V., LS Collier, MP Scott and CJ Tabin (1999). Vertebrate homologs of Drosophila Suppressor of fused interact with the Gli family of transcriptional regulators. Dev Biol 212: 323–336.

25. Jones-Villeneuve EM, MW McBurney, KA Rogers and VI Kalnins (1982). Retinoic acid induces embryonal carcinoma cells to differentiate into neurons and glial cells. J Cell Biol 94: 253–262.

26. Dunaeva M, P Michelson, P Kogerman and R Toftgard (2003). Characterization of the physical interaction of Gli proteins with SUFU proteins. J Biol Chem 278: 5116–5122.

27. Noguchi H, JG Castillo, K Nakashima and SJ Pleasure (2019). Suppressor of fused controls perinatal expansion and quiescence of future dentate adult neural stem cells. Elife 8: 1–21.

28. Hahn H, C Wicking, PG Zaphiropoulos, MR Gailani, S Shanley, A Chidambaram, I Vorechovsky, E Holmberg, AB Unden, S Gillies, K Negus, I Smyth, C Pressman, DJ Leffell, B Gerrard, AM Goldstein, M Dean, R Toftgard, G Chenevix-Trench, B Wainwright and AE Bale (1996). Mutations of the human homolog of drosophila patched in the nevoid basal cell carcinoma syndrome. Cell 85: 841–851.

29. McBurney M (1993). P19 embryonal carcinoma cells. Int J Dev Biol 140: 135–140.

30. Tan Y, Z Xie, M Ding, Z Wang, Q Yu, L Meng, H Zhu, X Huang, L Yu, X Meng and Y Chen (2010). Increased levels of FoxA1 transcription factor in pluripotent P19 embryonal carcinoma cells stimulate neural differentiation. Stem Cells Dev 19: 1365–1374.

31. St-Jacques B, M Hammerschmidt and AP McMahon (1999). Indian hedgehog signaling regulates proliferation and differentiation of chondrocytes and is essential for bone formation. Genes Dev 13: 2072–2086.

32. Bechtold TE, E Koyama, N Kurio, HD Nah, C Saunders and PC Billings (2019). The roles of Indian hedgehog signaling in TMJ formation. Int J Mol Sci 20:.

33. Bashamboo A and K McElreavey (2015). Human sex-determination and disorders of sex-development (DSD). Semin Cell Dev Biol 45: 77–83.

34. Bitgood MJ, L Shen and AP McMahon (1996). Sertoli cell signaling by Desert hedgehog regulates the male germline. Curr Biol 6: 298–304.

35. Clark AM, KK Garland and LD Russell (2000). Desert hedgehog (Dhh) gene is required in the mouse testis for formation of adult-type Leydig cells and normal development of peritubular cells and seminiferous tubules. Biol Reprod 63: 1825–1838.

36. Wu SM, ABH Choo, MGS Yap and KKK Chan (2010). Role of Sonic hedgehog signaling and the expression of its components in human embryonic stem cells. Stem Cell Res 4: 38–49.

37. Tang K, J Yang, X Gao, C Wang, L Liu, H Kitani, T Atsumi and N Jing (2002). Wnt-1 promotes neuronal differentiation and inhibits gliogenesis in P19 cells. Biochem Biophys Res Commun 293: 167–173.

38. Jing XT, HT Wu, Y Wu, X Ma, SH Liu, YR Wu, XF Ding, XZ Peng, BQ Qiang, JG Yuan, WH Fan and M Fan (2009). DIXDC1 promotes retinoic acid-induced neuronal differentiation and inhibits gliogenesis in P19 cells. Cell Mol Neurobiol 29: 55–67.

39. Chen W, X Caihong, B Wei, L Li, W Lin, Y-G Chen, S-L Ang and N Jing (2006). Cell aggregation-induced FGF8 elevation Is essential for P19 cell neural differentiation. Mol Biol Cell 17: 3075–3084.

40. Fraichard A, O Chassande, G Bilbaut, C Dehy, P Savatier and J Samarut (1995). In vitro differentiation of embryonic stem cells into glia and neutrons. J Cell Sci 108: 3181–3188.

41. Mak SK, YA Huang, S Iranmanesh, M Vangipuram, R Sundararajan, L Nguyen, JW Langston and B Schüle (2012). Small molecules greatly improve conversion of human-induced pluripotent stem cells to the neuronal lineage. Stem Cells Int 2012:.

42. Wu CY, D Whye, RW Mason and W Wang (2012). Efficient differentiation of mouse embryonic stem cells into motor neurons. J Vis Exp 1–5.

43. Okada Y, T Shimazaki, G Sobue and H Okano (2004). Retinoic-acid-concentration-dependent acquisition of neural cell identity during in vitro differentiation of mouse embryonic stem cells. Dev Biol 275: 124–142.

44. Li S, J Zheng, L Chai, M Lin, R Zeng, J Lu and J Bian (2019). Rapid and Efficient Differentiation of Rodent Neural Stem Cells into Oligodendrocyte Progenitor Cells. Dev Neurosci 41: 79–93.

45. Yue S, Y Chen and SY Cheng (2009). Hedgehog signaling promotes the degradation of tumor suppressor Sufu through the ubiquitin-proteasome pathway. Oncogene 28: 492–499.

46. Takenaka K, Y Kise and H Miki (2007). GSK3β positively regulates Hedgehog signaling through Sufu in mammalian cells. Biochem Biophys Res Commun 353: 501–508.

47. Infante P, R Faedda, F Bernardi, F Bufalieri, LL Severini, R Alfonsi, D Mazzà, M Siler, S Coni, A Po, M Petroni, E Ferretti, M Mori, E De Smaele, G Canettieri, C Capalbo, M Maroder, I Screpanti, M Kool, SM Pfister, D Guardavaccaro, A Gulino and L Di Marcotullio (2018). Itch/β-Arrestin2-dependent non-proteolytic ubiquitylation of SuFu controls Hedgehog signalling and medulloblastoma tumorigenesis. Nat Commun 9: 1–17.

48. Chen Y, S Yue, L Xie, XH Pu, T Jin and SY Cheng (2011). Dual phosphorylation of suppressor of fused (Sufu) by PKA and GSK3β regulates its stability and localization in the primary cilium. J Biol Chem 286: 13502–13511.

49. Wang Y, Y Li, G Hu, X Huang, H Rao, X Xiong, Z Luo, Q Lu and S Luo (2016). Nek2A phosphorylates and stabilizes SuFu: A new strategy of Gli2/Hedgehog signaling regulatory mechanism. Cell Signal 28: 1304–1313.

50. Chen MH, CW Wilson, YJ Li, KK Lo Law, CS Lu, R Gacayan, X Zhang, CC Hui and PT Chuang (2009). Cilium-independent regulation of Gli protein function by Sufu in Hedgehog signaling is evolutionarily conserved. Genes Dev 23: 1910–1928.

51. Hoelzl MA, K Heby-Henricson, G Bilousova, B Rozell, R V Kuiper, M Kasper, R Toftgård and S Teglund (2015). Suppressor of Fused plays an important role in regulating mesodermal differentiation of murine embryonic stem cells In vivo. Stem Cells Dev 24: 2547–2560.

52. Hoelzl MA, K Heby-Henricson, M Gerling, JM Dias, R V. Kuiper, C Trünkle, Å Bergström, J Ericson, R Toftgård and S Teglund (2017). Differential requirement of SUFU in tissue development discovered in a hypomorphic mouse model. Dev Biol 429: 132–146.

53. Kise Y, A Morinaka, S Teglund and H Miki (2009). Sufu recruits GSK3β for efficient processing of Gli3. Biochem Biophys Res Commun 387: 569–574.

54. Yabut OR, G Fernandez, K Yoon, SJ Pleasure, OR Yabut, G Fernandez, T Huynh, K Yoon and SJ Pleasure (2015). Suppressor of Fused Is critical for maintenance of neuronal progenitor identity during corticogenesis. Cell Rep 12: 2021–2034.

55. Jiwani T, JJ Kim and ND Rosenblum (2020). Suppressor of fused controls cerebellum granule cell proliferation by suppressing Fgf8 and spatially regulating Gli proteins. Development 147: 1–17.

