## Supplemental Table and Figures for "Suppressor of Fused regulation of Hedgehog Signaling is Required for Proper Astrocyte Differentiation"

**Supplemental Table 1** – Primers used for gene amplifications and CRISPR-Cas9

| CRISPR-Cas9 | | |
|---|---|---|
| *Sufu* gRNA | GGCTGATAACTGACATGCGG | |
| | Forward Primer (5′ → 3′) | Reverse Primer (5′ → 3′) |
| *Sufu* gRNA | CACCGGGCTGATAACTGACATGCGG | AAACCCGCATGTCAGTTATCAGCCC |
| *Sufu* PCR | CTCCATCCCACCTGTAGAGTTC | AGCAAGGTTTTCCTCACTCAAG |
| *U6* | GGGCAGGAAGAGGGCCTAT | |
| **qRT-PCR** | | |
| **Gene** | Forward Primer (5′ → 3′) | Reverse Primer (5′ → 3′) |
| *L14* | GGGTGGCCTACATTTCCTTCG | GAGTACAGGGTCCATCCACTAAA |
| *Shh* | AAAGCTGACCCCTTTAGCCTA | TTCGGAGTTTCTTGTGATCTTCC |
| *Ihh* | GACTCATTGCCTCCCAGAACTG | CCAGGTAGTAGGGTCACATTGC |
| *Dhh* | ACCCCGACATAATCTTCAAGGAT | GTACTCCGGGCCACATGTTC |
| *Smo* | CCTGACTTTCTGCGTTGC | GGTTCTGACACTGAATCCG |
| *Ptch1* | AAAGAACTGCGGCAAGTTTTTG | CTTCTCCTATCTTCTGACGGGT |
| *Ptch2* | CTCCGCACCTCATATCCTAGC | TCCCAGGAAGAGCACTTTGC |
| *Sufu* | CGGACCCCTTGGACTATGTTA | CTTCAGACGAAACGTCAACTCA |
| *Gli1* | GGAAGTCCTATTCACGCCTTGA | CAACCTTCTTGCTCACACATGTAAG |
| *Gli2* | TACCTCAACCCTGTGGATGC | CTACCAGCGAGTTGGGAGAG |
| *Gli3* | CTGTCGGCTTAGGATCTGTTG | GCTCTTCAGCAAGTGGTTCC |
| *Ascl1* | ACTTGAACTCTATGGCGGGTT | CCAGTTGGTAAAGTCCAGCAG |
| *NeuroD1* | GCATGCACGGGCTGAACGC | GGGATGCACCGGGAAGGAAG |
| *NeuroG1* | CCAGCGACACTGAGTCCTG | CGGGCCATAGGTGAAGTCTT |
| *Chicken Sufu* | ACGGGCAAGGAATCTTGGAG | ACTCTCTCTTGCAGATGCGG |

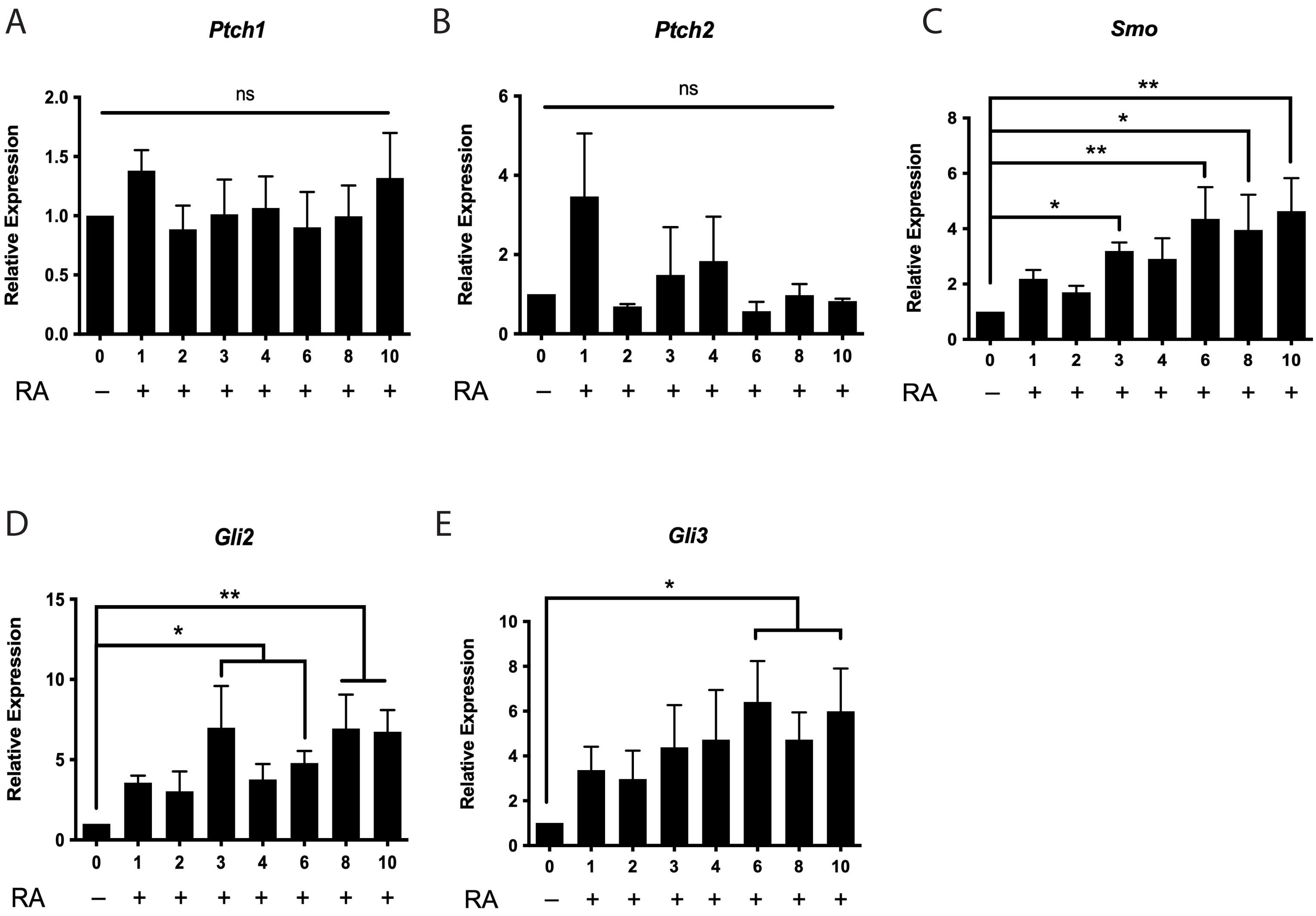


**Supplemental Figure 1 – Expression of Hh pathway components during RA-induced differentiation.** Relative expression of **(A)** *Ptch1*, **(B)** *Ptch2*, **(C)** *Smo*, **(D)** *Gli1*, **(E)** *Gli2* and **(F)** *Gli3*. N=3. Bars represent mean values plus/minus s.e.m. *$P<0.05$, **$P<0.01$.

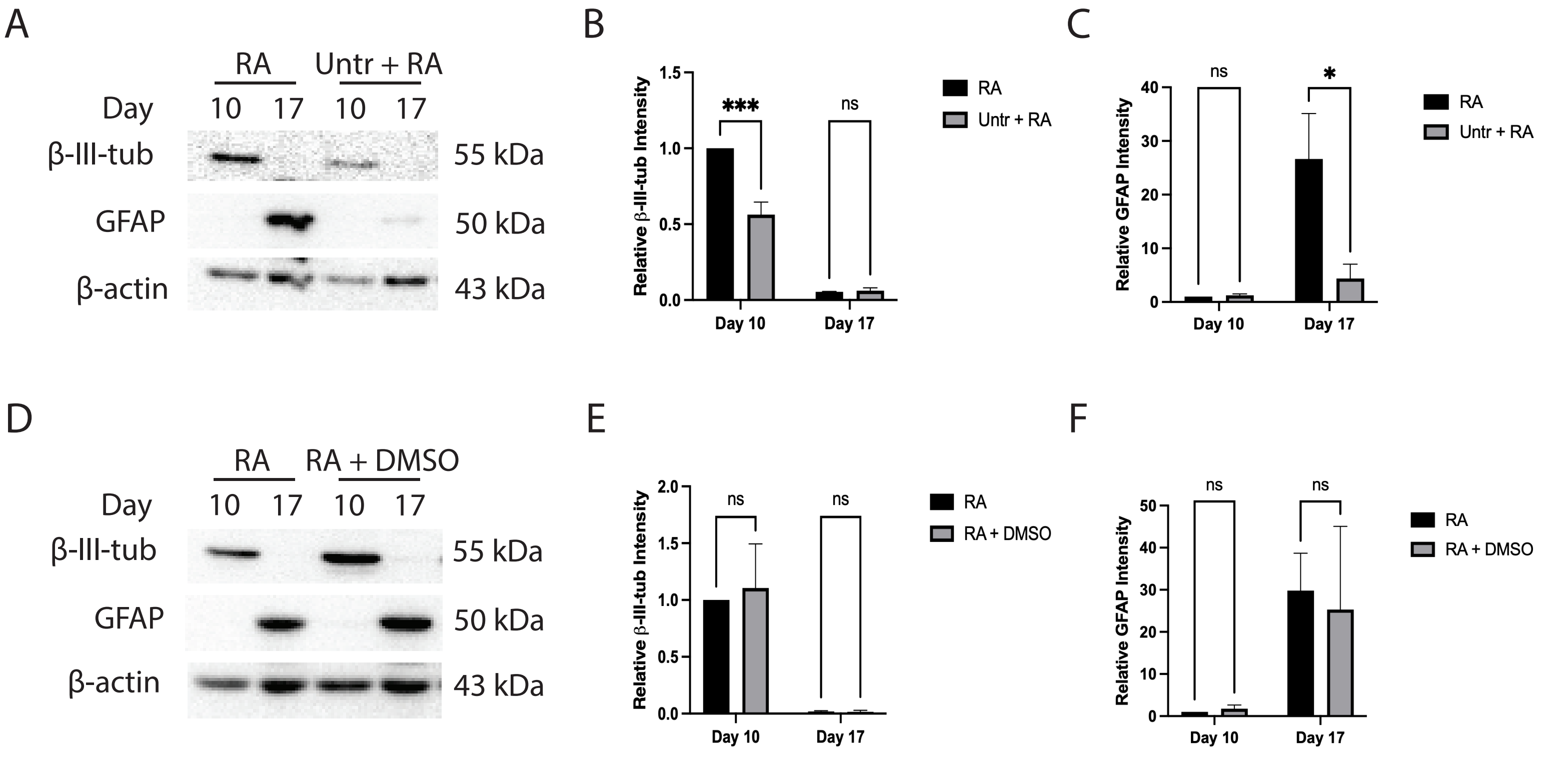


**Supplemental Figure 2 – Untreated embryoid bodies plus RA is not sufficient to induce differentiation**. **(A)** Immunoblot of beta-III-tubulin and GFAP on days 10 and 17 of cells induced to differentiate in the presence of RA alone, or untreated cells in embryoid bodies plus RA treatment on day 4. **(B)** Densitometry of beta-III-tubulin and **C)** of GFAP from immunoblot in A. **(D)** Immunoblot of beta-III-tubulin and GFAP on days 10 and 17 of cells induced to differentiation in the presence of RA alone or RA plus vehicle control of Cyclopmaine, DMSO. **(E)** Densitometry of beta-III-tubulin and **(F)** GFAP from immunoblot in D. N=3. Bars represent mean values plus/ minus s.e.m.

**P<0.05, ***P<0.001.*

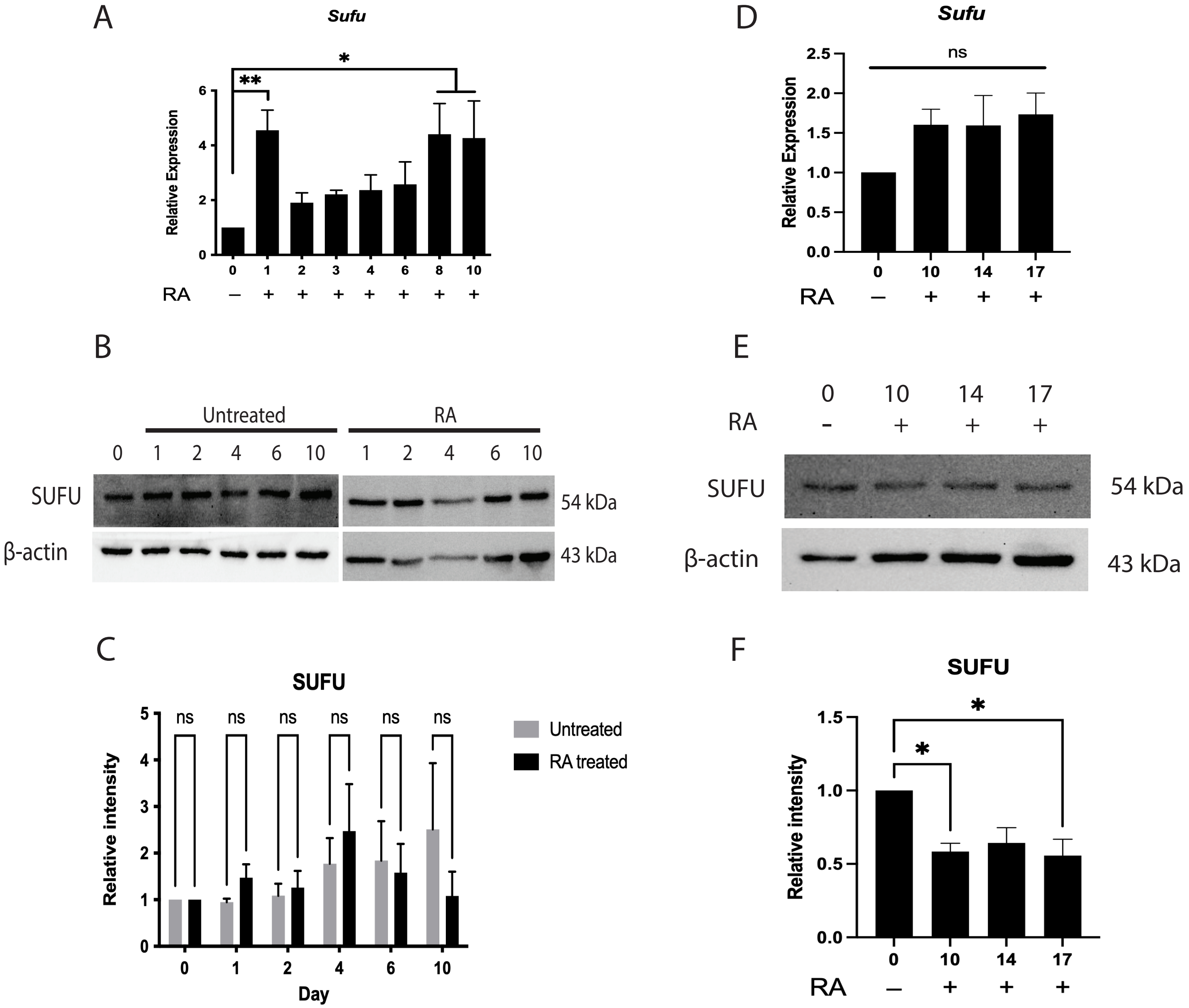


**Supplemental Figure 3 – SUFU levels during RA-induced differentiation. (A)** Expression of *Sufu* on days 0-10. **(B)** Immunoblot of SUFU protein on days 0-10 and **(C)** densitometric analysis of the immunoblot. **(D)** Expression of *Sufu* on days 0-17, **(E)** immunoblot analysis of the SUFU protein at these time points, and **(F)** densitometric analysis of the immunoblot in E. N=3. Bars represent mean values plus/minus s.e.m.*$P<0.05$, **$P<0.01$.

A

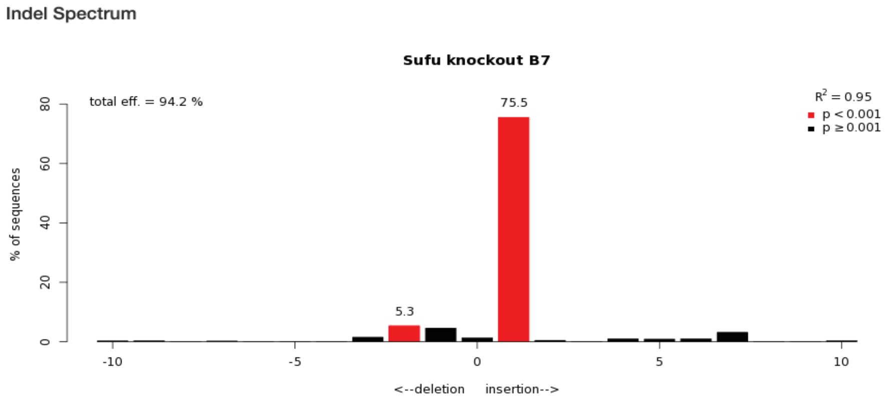


B

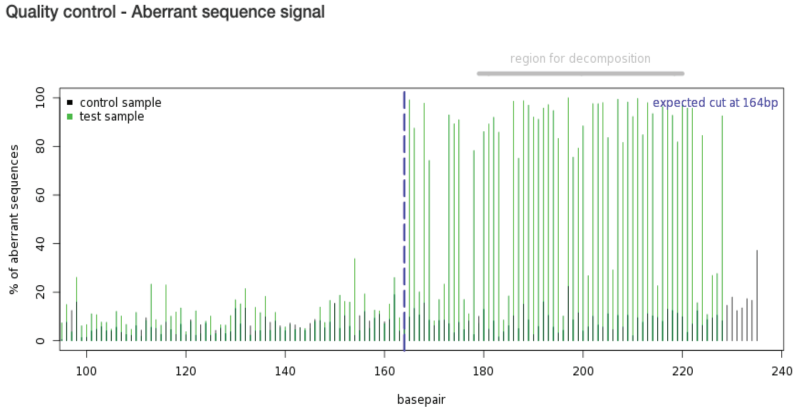


**Remarks**

Default settings:
alignment window = 100 - 154
decomposition window = 179 - 220
indel size = 10
p threshold = 0.001

C

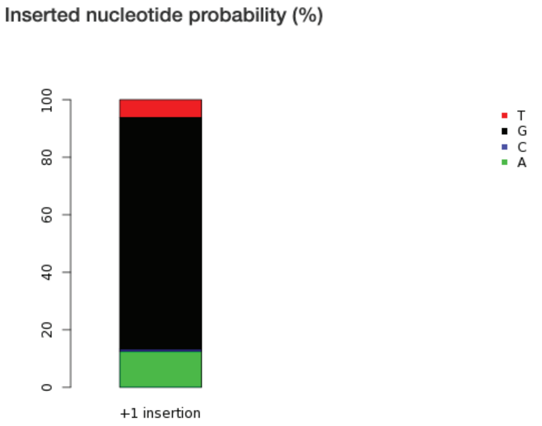


| A | C | G | T |
|---|---|---|---|
| 12.5 | 0.7 | 80.6 | 6.2 |

**Supplemental Figure 4 – TIDE analysis of Sufu-/- clone (KO#2). (A)** Indel spectrum analysis showing predicted mutant allele after sequence alignment. Bars represent percent of sequences identified during analysis, with red bars being statistically significant. **(B)** Aberrant sequence analysis after sequence alignment. Black lines represent wildtype sequence and green lines represent alignment of mutant sequence. Dashed line represents the predicted cut site. Taller green compared to black lines indicate a likely mutation. **(C)** Inserted nucleotide probability score after Indel spectrum analysis.

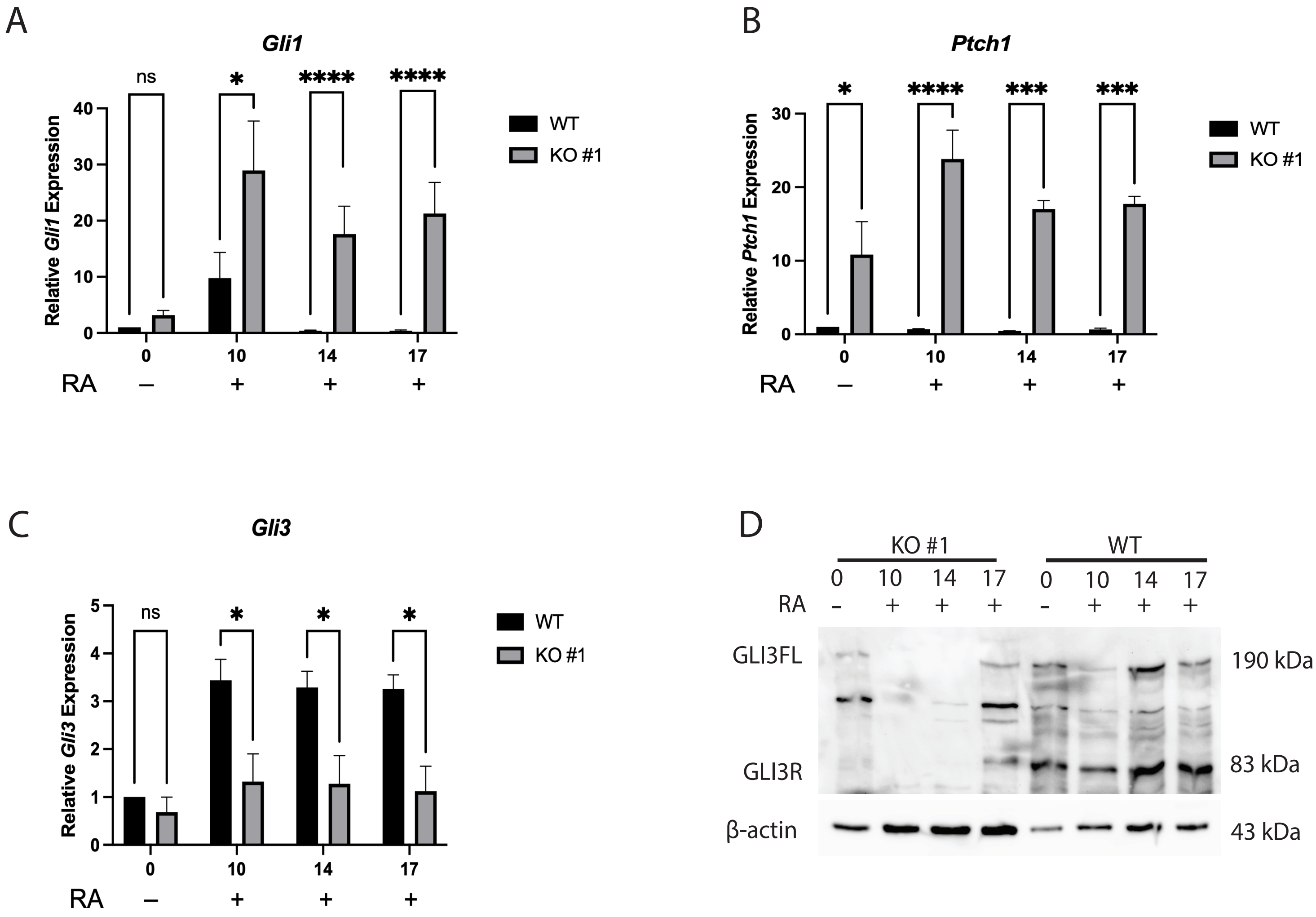


**Supplemental Figure 5 –*Sufu-/-* maintains activate Hh signaling and causes the loss/reduction of Gli3 in KO #1.** Expression of Hh target genes **(A)** *Gli1* and **(B)** *Ptch1* and **(C)** *Gli3* encoding transcription factor in wildtype (WT) and *Sufu-/-* KO #1 cells on days 0-17. **(D)** Immunoblot of GLI3 full length (GLI3FL) and GLI3 repressor (GLI3R) in WT and *Sufu-/-* KO #1 cells on days 0-17. N=3. Bars represent mean values plus/minus s.e.m. **P<0.05, **P<0.01, ***P<0.001, ****P<0.0001.*

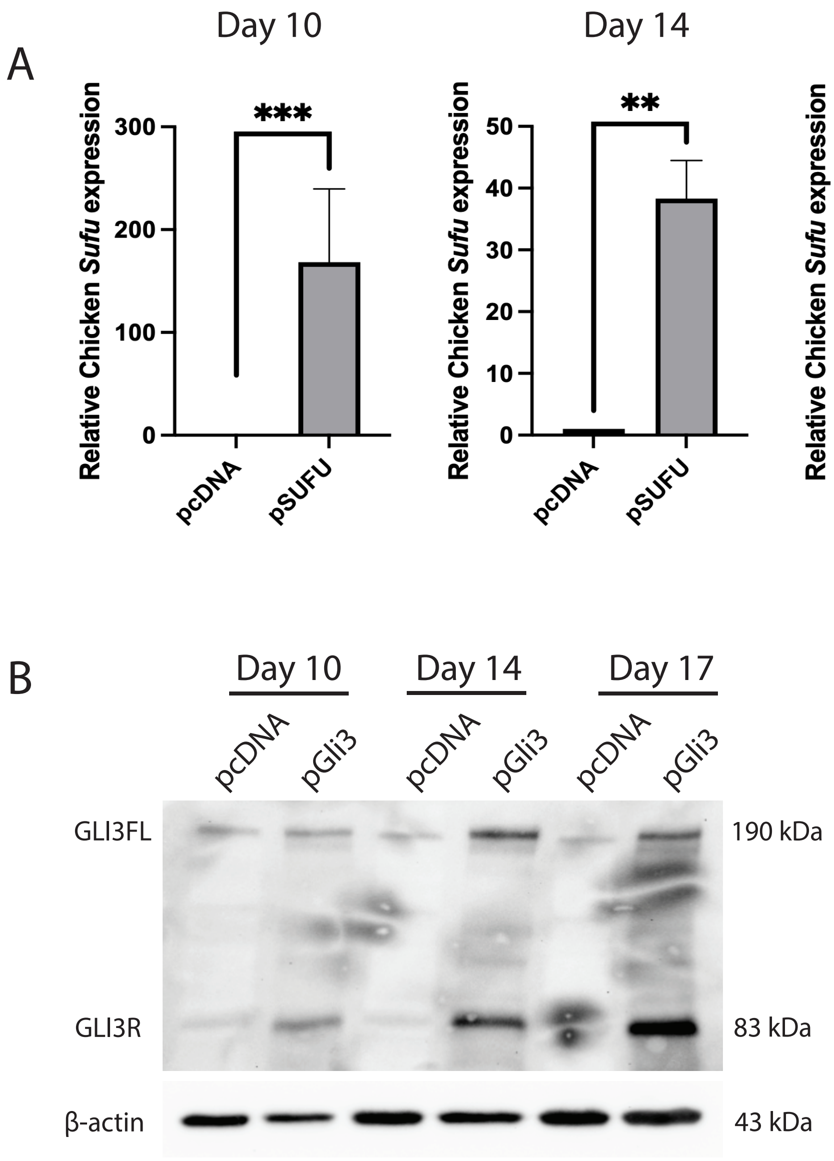


**Supplemental Figure 6 - pSUFU and pGli3 overexpression induces Chicken *Sufu* expression and rescued detection of GLI3. (A)** Chicken *Sufu* expression was detected using Chicken-specific RT-qPCR primers at days 10, 14 and 17 of RA treatment in pcDNA and pSUFU stably transfected Sufu-/- cells. **(B)** GLI3 fullength (GLI3FL) and repressor (GLI3R) was detected at days 10, 14 and 17 of RA in pGLI3 stably transfected Sufu-/- cells. N=3. Bars represent mean values +/- s.e.m. *$**P<0.01$, $***P<0.001$*.
